# Turtle IgD2 preserves an ancestral IgXA-derived XA3–XA4 module in a duplicated and locally remodeled IgD2–IgY constant-region array

**DOI:** 10.64898/2026.07.17.739153

**Authors:** Francisco Gambón-Deza

**Author notes:** Correspondence should be addressed to Francisco Gambón-Deza.

## Abstract

A second IgD gene, IgD2, was first described in the leopard gecko as a hybrid constant-region gene containing IgD-derived exons and terminal IgA-like exons. Related IgD2 architectures have subsequently been recognized in turtles and crocodilians. Here we analyzed Testudines constant-region annotations to determine whether turtle IgD2 represents an intact ancient paralogue or a locally remodeled mosaic gene. Across representative turtle genomes, mixed loci frequently contained upstream D1–D4 exons followed by 3_XA and 4_XA. A direct comparison of paired turtle XA3–XA4 modules with amphibian IgXA and IgM references showed that all turtle modules were closer to amphibian IgXA than to either amphibian or turtle IgM. Upstream D exons, especially D1–D3, instead showed strong local similarity to canonical IgD exons within the same species. A conservative nucleotide tract-permutation test confirmed clustered conversion-like tracts in a subset of D1 and D2 exons, whereas D3 retained local affinity without significant tract clustering. IgY analysis identified 76 complete Y1–Y4 blocks; most were genomically proximal to XA-bearing IgD2 loci on the opposite strand, and contig-restricted randomization confirmed that this association was non-random. All six amino-acid distance-matrix comparisons among Y1–Y4 were significantly correlated, supporting a coupled IgY history without a D-like domain-specific rupture. Independent reannotation recovered 13 complete, one-to-one IgD2/XA–IgY pairs in *Chelonia mydas*; a thymus transcript encoded the complete D1–D4/XA3– XA4 architecture. *Dermochelys coriacea* provided three further complete pairs. In *Mauremys reevesii*, 14 oppositely oriented modules form an array marked by segmental duplication, strand switches, and exon loss. These results support an ancestral paired IgD2–IgY architecture that expanded by segmental duplication to generate the present germline array, while a subset of IgD2 D1 and D2 exons continued to undergo recent intra-species exchange with canonical IgD-derived material. The genomic mechanism can be reconstructed, but the biological significance of maintaining this unusual opposite-strand association remains unknown.

## 1 Introduction

The immunoglobulin heavy-chain locus is among the most dynamic regions of vertebrate immune genomes. Constant-region genes can be expanded, contracted, rearranged, or homogenized by gene conversion, producing lineage-specific antibody isotypes and domain architectures. IgM is the ancestral secreted antibody class of jawed vertebrates, whereas IgD shows exceptional structural variability across vertebrate lineages.

IgD2 was originally described in the leopard gecko, *Eublepharis macularius*, as a second IgD gene generated by duplication of an older immunoglobulin gene and recombination with an IgA-like gene [1, 2]. This observation established that reptile IgD evolution could produce hybrid constant-region genes rather than only canonical long IgD genes. Reptile heavy-chain loci were subsequently characterized in *Anolis carolinensis*, snakes, turtles, and crocodilians, revealing conserved IgM and IgD components but also lineage-specific IgY, IgA, and IgD2-like complexity [3, 4, 5, 6, 7]. In particular, IgD2-like genes were reported in turtles and crocodilians, with duplications and domain losses in painted and soft-shelled turtles [5, 6, 10]. Thus, Testudines provide a critical system for asking whether IgD2 is an old amniote architecture or a repeatedly reconstructed modular gene. In amphibians, previous work described IgD genes related to ancestral amniote IgD and two IgA/X isotypes [8]. A subsequent reconstruction of amphibian heavy-chain loci resolved IgXA as a four-exon mosaic immunoglobulin: XA1 and XA2 show IgY-like ancestry, XA3 has an intermediate history, and XA4 retains the clearest IgM-like signal [9]. In turtles, preliminary annotation of heavy-chain constant-region exons revealed a recurrent non-canonical architecture: IgD-like exons upstream of two unusual exons, named 3_XA and 4_XA in our annotation system.

The biological interpretation of this architecture is not straightforward. The turtle 3_XA and 4_XA exons are expected, under the amphibian IgXA model, to represent the retained XA3– XA4 part of an older IgXA module rather than newly evolved IgD exons. The open question is therefore not whether XA exons are IgD D exons, but how this ancestral XA3–XA4 module became embedded in turtle IgD-like constant-region arrays. If the whole gene were a true ancient IgD2 paralogue, the upstream D exons and the XA3–XA4 module should share a coherent orthologous history across turtle families. If the structure arose independently within each lineage, the XA-bearing genes should group with local canonical IgD copies. A third possibility is mosaic evolution: the XA3–XA4 module may have a deep IgXA origin, while the upstream D exons may be recruited repeatedly from local IgD-like constant-region material.

Here we test these alternatives using protein and nucleotide sequences from annotated turtle constant-region exons. We focus on three questions: (i) do turtle 3_XA and 4_XA behave as a conserved IgXA-derived module distinct from IgD D exons; (ii) do the upstream D exons of XA-bearing genes have a single trunk origin comparable to canonical 11-domain IgD; and (iii) is the final architecture better explained by orthology, local duplication, or gene conversion?

## 2 Materials and Methods

### 2.1 Input data and exon calls

Analyses were performed on locally generated Testudines heavy-chain constant-region exon annotations derived from public turtle genome assemblies. For each species, the chs directory was parsed for amino-acid exon sequences (aexones.fasta), nucleotide exon sequences (nexones.fasta), and annotation assignments (aexones_asignados.txt). Candidate exons were retained when their assignment score was at least 0.90. Overlapping calls of the same exon class on the same species, contig, and strand were reduced to the best-scoring, longest representative.

The VGP primary assembly of *Chelonia mydas* (GCA_015237465.2) was reprocessed with the same CHS detector used for the final Testudines annotations, and high-confidence calls on CM026910.2 were reconstructed de novo. The *Dermochelys coriacea* comparison used GCA_009764565.4 and its final CHS calls on CM019947.2. A previously assembled *C. mydas* thymus open reading frame from SRA run SRR14839285 was inspected as independent transcript evidence for the D1–D4/XA3–XA4 architecture.

A submission-oriented sampling table listing species, genome assembly identifiers, CHS exon counts, representative IgM contigs, representative mixed-locus contigs, and complete IgY block counts is provided as Supplementary Table S1. One-to-one array counts and IgY domain-congruence statistics are provided as Supplementary Tables S4 and S5, respectively.

### 2.2 Definition of mixed IgD2-like loci

Mixed loci were defined operationally as genomic blocks containing one or more XA exons and upstream exons annotated as IgD-like D exons. Blocks were reconstructed by ordering exons on each species, contig, and strand in transcriptional direction. For minus-strand genes, genomic coordinates were reversed before block construction. A new block was started at a new D1 exon, at a large inter-exon gap, or when the D-exon order reset. For each XA-bearing block, only D exons located upstream of the first XA exon were considered part of the upstream D scaffold.

Canonical IgD references were selected per species as the non-XA block with the largest number of distinct D exons, prioritizing blocks approaching the 11-domain IgD architecture. Representative mixed blocks were also chosen per species, prioritizing complete D1–D4 + 3_XA + 4_XA architectures.

### 2.3 Sequence alignment and distance analyses

Protein and nucleotide sequences were aligned with MAFFT v7.526 using the --auto strategy [11]. Pairwise amino-acid and nucleotide p-distances were computed after excluding gaps and ambiguous residues. For nucleotide alignments, Jukes–Cantor 1969 corrected distances were also calculated [16]. Neighbor-joining trees were generated from p-distance matrices for exploratory topology inspection [15]. Conversion-like cases were defined as mixed-locus upstream D exons whose nearest canonical IgD exon was closer than any mixed-locus exon from another species.

For D1–D3 nucleotide alignments, local conversion was additionally tested with a tract-permutation procedure. Each mixed-locus exon was compared with the best same-species canonical IgD exon and with the nearest canonical IgD exon across all species. The longest un-gapped identical tract between the mixed exon and the same-species canonical exon was recorded. Significance was estimated by 5000 permutations of the observed match/mismatch vector, preserving the total number of identical and different sites while randomizing their order. Permutation *p*-values were corrected within each exon class using the Benjamini–Hochberg procedure. A strict conversion-supported case required: (i) the nearest canonical exon to be from the same species, (ii) tract *q ≤* 0.10, and (iii) same-species canonical–mixed p-distance *≤* 0.12.

### 2.4 Reconstruction of IgM amino-acid sequences

To provide a constant-region reference tree for the sampled turtles, IgM blocks were reconstructed from the last/*/CHS annotation outputs generated for each genome. Exons annotated as M1–M4 were ordered by species, genome, contig, and strand, using transcriptional order for minus-strand loci. Complete M1–M4 blocks were retained when all four distinct M exons were present in a single local block. For species with more than one complete block or assembly, one representative was selected by the highest mean annotation probability and then by total amino-acid length. The selected M1–M4 amino-acid sequences were concatenated, labeled as IgM-followed by the species name, aligned with MAFFT, and used to generate an amino-acid p-distance neighbor-joining tree. Node support was estimated by non-parametric bootstrap resampling of alignment columns with 1000 replicates; bootstrap values are shown as percentages on the internal branches of the IgM tree.

### 2.5 IgY block reconstruction and syntenic analysis

IgY blocks were reconstructed from Y1–Y4 exon calls in the same last/*/CHS outputs. Complete IgY genes were defined as local blocks containing Y1, Y2, Y3, and Y4 in transcriptional order. Complete Y1–Y4 amino-acid sequences were concatenated, aligned with MAFFT, and analyzed by pairwise evolutionary p-distance and nearest-neighbor relationships. A neighbor-joining tree was also reconstructed from the complete Y1–Y4 alignment, with node support estimated by 1000 non-parametric bootstrap replicates. Because several species contain multiple complete IgY copies, tree labels were formatted as IgY-species-copy. For each IgY block, the nearest XA, D, and M exon calls were recorded, including physical genomic separation, contig, and strand. IgY blocks within 100 kb of an XA exon were considered syntenically associated with the IgD2/XA array; the strand relationship was used to test the observation that IgD2/XA genes and repeated IgY genes commonly lie on complementary strands.

The genomic proximity of complete IgY blocks to XA-bearing IgD2 loci was tested by randomization within contigs. The test was restricted to complete IgY blocks that had high-confidence XA3 or XA4 calls on the same contig. For each evaluable IgY block, the observed distance and strand relationship to the nearest XA exon were recorded. In each of 20,000 permutations, the IgY block length and contig were preserved, but the block start was randomized uniformly within the local span covered by IgY and XA annotations on that contig. The primary statistic was the number of complete IgY blocks located within 100 kb of an opposite-strand XA exon. Empirical *p*-values were calculated as (*r* + 1)*/*(*n* + 1), where *r* is the number of randomizations with a statistic at least as extreme as the observed value and *n* is the number of randomizations.

To test whether IgY domains retain a coupled history, the same 76 complete IgY blocks were aligned separately for Y1, Y2, Y3, and Y4. Amino-acid p-distance matrices were calculated for each domain, and Spearman correlations were computed for all six pairs of upper-triangular distance matrices. Significance was assessed with 10,000 label permutations that preserved each matrix but randomized the correspondence among genes. For each gene and domain, the nearest sequence was also recorded; the fraction of genes with the same nearest neighbor in each domain pair was used as an intuitive measure of local topological agreement.

For the comparative physical test, complete IgY blocks were paired one-to-one with the nearest opposite-strand XA-bearing IgD2 block on the same contig when their interval distance was no greater than 100 kb. The *C. mydas* and *D. coriacea* array intervals, with 20 kb margins, were aligned against themselves with BLASTN. Alignments of at least 700 bp and at least 70% nucleotide identity were retained, excluding the trivial forward diagonal, and direct and inverted matches were plotted separately.

### 2.6 Module-level analysis of the *Mauremys reevesii* IgD2–IgY array

The expanded *Mauremys reevesii* constant-region locus on NC_052635.1 was re-examined directly from the last/CHS/result-chs.tsv output. Accepted exons that were present in the final annotation but absent from downstream block tables were restored to a curated local table, and two IgY3 candidates located in IgY block gaps were retained as putative calls for architectural inspection. IgD2/XA and IgY blocks were paired when they were located on opposite strands and separated by no more than 30 kb. Each pair was treated as a candidate duplicated module and represented by the ordered amino-acid components D1–D4, XA3–XA4, and Y1–Y4, using gaps for missing exons.

To test physical duplication directly, the NC_052635.1 interval spanning all curated pairs plus 35 kb margins on each side was extracted from the genome assembly and aligned against itself. The extracted interval covered NC_052635.1:3,667,666–4,315,997. Self-alignments were generated with BLASTN using local nucleotide alignments of at least 700 bp and at least 70% identity; same-orientation and inverted alignments were plotted separately. Pair-level overlap counts were then calculated by intersecting self-alignment segments with each curated IgD2/XA–IgY module.

Recovered exons added during the *Mauremys reevesii* curation were validated against the genomic sequence. For each rescued or putative exon, the transcript-oriented nucleotide sequence was extracted and all three coding phases were screened for stop-free open reading frames. Exon length was compared with high-confidence accepted exons of the same label where available. Because candidate coordinates may not exactly match splice junctions, canonical splice motifs were scored both at the annotated boundary and within a *±*12 nt window in transcript orientation. Validation calls were classified as strong when ORF, expected length, and nearby splice signals were all supported; as coding/splice-supported without length reference when no local length reference was available; or as questionable when a stop-free ORF was not recovered.

To compare the evolutionary histories of different module components, amino-acid trees were reconstructed for four concatenations from the same paired units: the full D1–D4/XA3–XA4/Y1– Y4 module, the potentially exchangeable D1–D3 scaffold, the IgY Y1–Y4 component, and the XA3–XA4 component. Each component set was aligned with MAFFT, pairwise amino-acid p-distances were calculated after excluding gaps, and neighbor-joining trees were midpoint-rooted. Node support was estimated with 500 bootstrap replicates. In addition, maximum-likelihood trees were reconstructed for the IgY Y1–Y4, D1–D3, and XA3–XA4 component alignments using IQ-TREE v3.1.2 [12]. ModelFinder was used to select the best-fit amino-acid model under BIC [13], and node support was estimated with 1000 ultrafast bootstrap replicates [14]. ML trees were midpoint-rooted for comparison with the NJ component trees.

### 2.7 Testing the retained IgXA3–IgXA4 signal

Because amphibian IgXA reconstruction identified an intermediate history for XA3 and the clearest IgM-like ancestry in XA4, we compared turtle 3_XA and 4_XA against D and M exon calls as an internal consistency test. High-confidence D, M, and XA exon calls were extracted for exon 3 and exon 4 classes. Protein alignments were generated separately for exon3_D/exon3_M/exon3_XA and exon4_D/exon4_M/exon4_XA. Mean amino-acid p-distances among D, M, and XA classes were then compared. These analyses were not treated as de novo proof of IgXA origin, but as a test of whether turtle XA3/XA4 retain the expected distinction from IgD and compatibility with the amphibian IgXA model.

### 2.8 Direct comparison with amphibian IgXA and IgM references

To distinguish inherited IgXA ancestry from a recent turtle-specific IgM duplication, we compared paired turtle XA3–XA4 modules with amphibian XA3–XA4 and IgM references. Amphibian constant-region sequences were taken from the reconstructed genome-based dataset used to define the mosaic origin of amphibian IgXA [9], which extends the earlier comparative amphibian IgA/X framework [8]. The comparison comprised 12 amphibian IgXA3–IgXA4 pairs and three amphibian IgM3–IgM4 pairs. Species, sequence accessions, strand, exon coordinates, annotation scores, and amino-acid lengths are listed in Supplementary Table S2. Pairs were retained only when both exons occurred in the same species, contig, and strand, in the expected transcriptional order, and within a 20 kb interval. Turtle XA3–XA4 modules were reconstructed from representative mixed loci using the same order rule. XA3 and XA4 amino-acid sequences were concatenated with a short spacer and aligned with amphibian IgXA3–IgXA4, amphibian IgM3–IgM4, and turtle IgM3–IgM4 references. Pairwise amino-acid p-distances and nearest-neighbor relationships were then calculated for all turtle XA3–XA4 modules.

## 3 Results

### 3.1 A reconstructed IgM tree defines the sampled Testudines framework

The IgM reconstruction identified 275 M exon calls distributed into 108 local M blocks. Seventeen blocks contained a complete M1–M4 structure, and after choosing one representative per species or assembly group, 15 species-level IgM amino-acid sequences were retained for phylogenetic reconstruction. These sequences were concatenated as M1–M4, aligned, and labeled as IgM-species in the resulting neighbor-joining tree with 1000 bootstrap replicates (Fig. 1).

**Figure 1:**
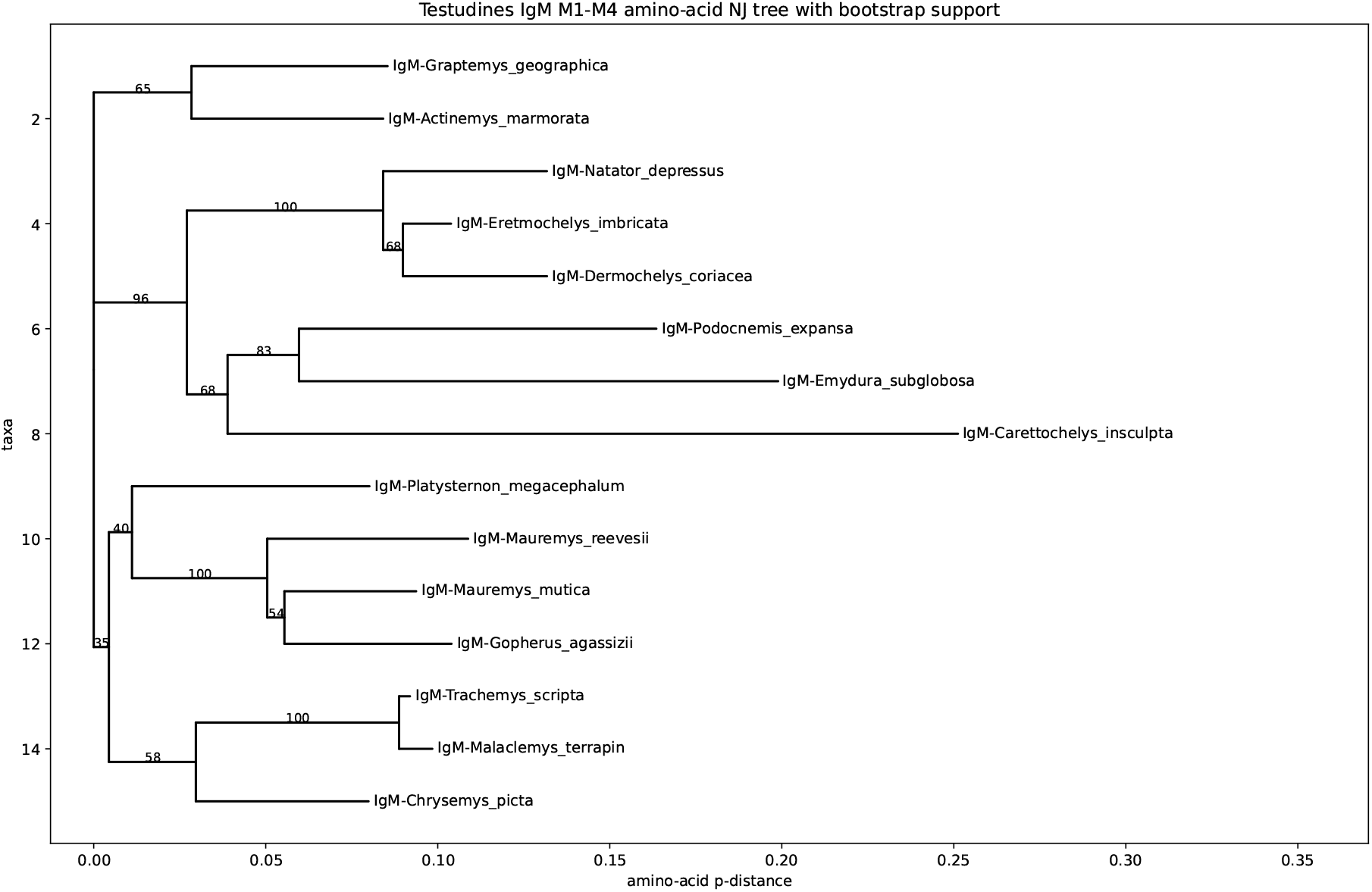
IgM phylogenetic framework for the sampled Testudines. Neighbor-joining tree reconstructed from concatenated M1–M4 amino-acid sequences. Labels identify each sequence as IgM-species; node values show support from 1000 non-parametric bootstrap replicates. The tree is used as an internal constant-region reference and is not intended as a formal species-tree estimate.

The tree provides an internal constant-region reference for the turtle dataset rather than a formal species-tree estimate. Several expected affinities are recovered with moderate to high bootstrap support, including clustering of marine turtles *Eretmochelys imbricata*, *Natator depressus*, and *Dermochelys coriacea*, and grouping of geoemydid *Mauremys* species. Other placements should be interpreted cautiously because the tree is based only on concatenated IgM constant-region exons and some deeper nodes have low bootstrap support. Nevertheless, it confirms that the IgM sequences reconstructed from the same annotation framework are suitable as comparative references for distinguishing IgM-like signal from IgXA-derived signal in the mixed loci.

### 3.2 XA-bearing genes are recurrent mixed loci rather than isolated exon calls

Representative mixed loci were detected across phylogenetically diverse turtles (Fig. 2; conceptual model in Supplementary Fig. S1). Complete or near-complete D1–D4 + 3_XA + 4_XA structures were present in *Chrysemys picta*, *Dermochelys coriacea*, *Eretmochelys imbricata*, *Graptemys geographica*, *Malaclemys terrapin*, *Mauremys mutica*, and *Mauremys reevesii*. *Gopherus agassizii* contained a partial D2–D3 + 3_XA + 4_XA block, whereas *Emydura subglobosa* showed an atypical D1–D2–D3–D9 + 3_XA + 4_XA architecture.

**Figure 2:**
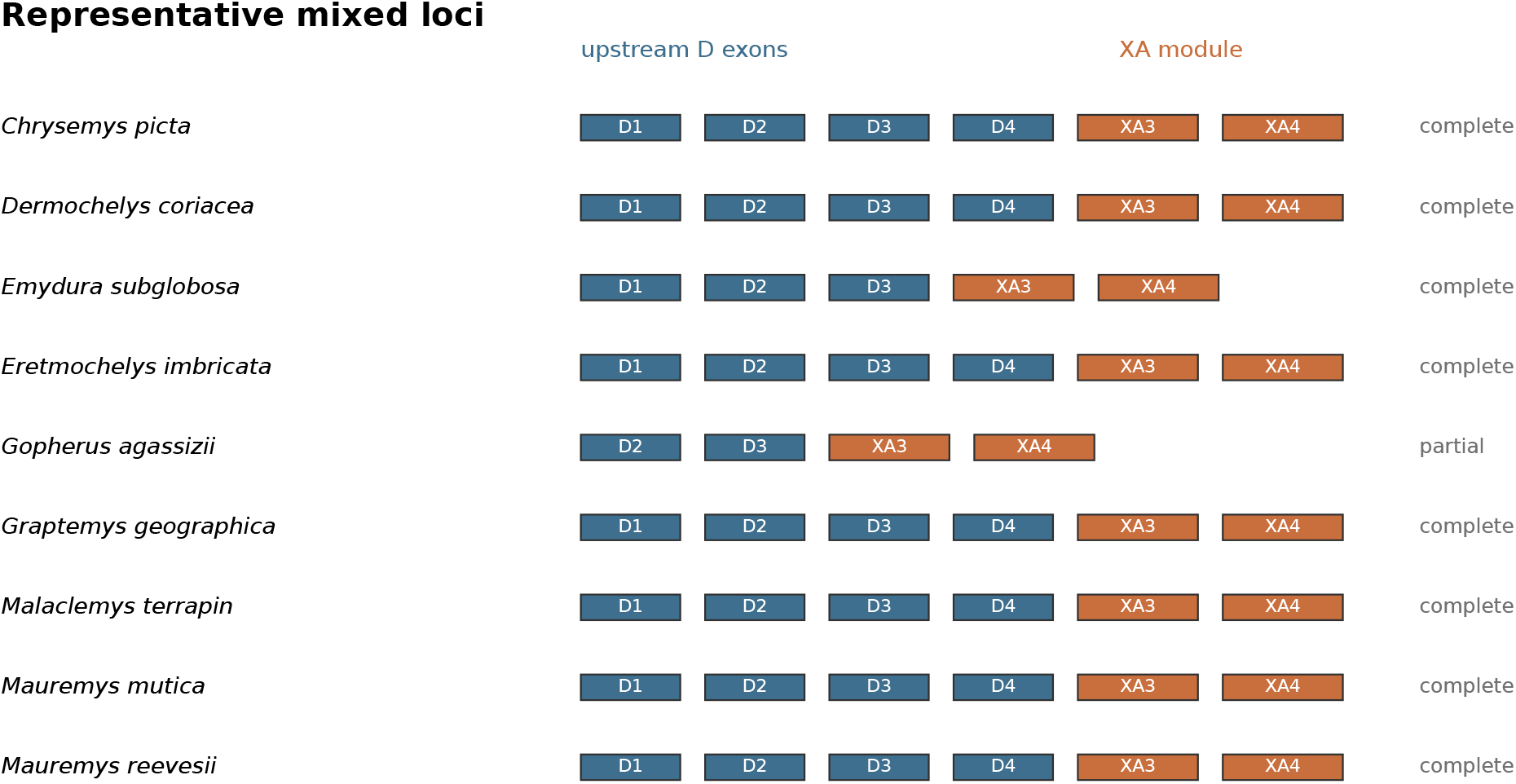
Representative mixed loci across Testudines. Each row shows one representative XA-bearing block per species. Blue boxes represent upstream IgD-like D exons and orange boxes represent XA exons. Most sampled species contain a D1–D4 + XA3–XA4 architecture, with partial or atypical structures in *Gopherus agassizii* and *Emydura subglobosa*.

This repeated syntenic arrangement indicates that XA exons are not sporadic annotation artifacts. Instead, they define a recurrent constant-region module embedded within IgD-like genomic neighborhoods.

### 3.3 Turtle XA3/XA4 are distinct from IgD and retain the expected IgXA signal

Protein distances showed that turtle XA3/XA4 exons are not close variants of IgD D exons. For exon 3, mean D–XA distance was 0.707, while XA–XA distance was 0.111. For exon 4, mean D–XA distance was 0.658, while XA–XA distance was 0.082. Thus, turtle XA exons form a coherent class separate from IgD D exons, as expected if they represent retained components of the older IgXA immunoglobulin.

Comparison with turtle IgM exons was consistent with the published amphibian IgXA model but did not need to independently establish it (Supplementary Fig. S2). For exon 4, M–XA distances were lower than D–XA distances (0.608 versus 0.658), and all 4_XA sequences had a nearest M exon closer than their nearest D exon when compared across classes. For exon 3, M–XA and D–XA distances were both high (0.699 and 0.707), and the nearest-neighbor signal was mixed.

We interpret these results as showing that turtle 3_XA and 4_XA are not IgD D exons and that 4_XA, in particular, retains the expected IgM-related signal of the ancestral IgXA module, whereas XA3 preserves the intermediate affinity described in amphibians.

### 3.4 Turtle XA3–XA4 is closer to amphibian IgXA than to IgM

The direct paired-module test supported homology between turtle XA3–XA4 and amphibian IgXA3–XA4. Mean amino-acid p-distance among turtle XA3–XA4 modules was 0.144. The mean distance between amphibian IgXA3–IgXA4 and turtle XA3–XA4 was 0.611, lower than the corresponding distances from amphibian IgM3–IgM4 to turtle XA3–XA4 (0.624) and from turtle IgM3–IgM4 to turtle XA3–XA4 (0.654) (Fig. 3). Most importantly, all nine turtle XA3–XA4 modules had an amphibian IgXA3–IgXA4 sequence closer than any IgM reference. The 4_XA exon retains the strongest IgM-like signal expected from the mosaic ancestry of IgXA, while the paired XA3–XA4 module behaves as IgXA rather than as an independent turtle IgM duplication.

**Figure 3:**
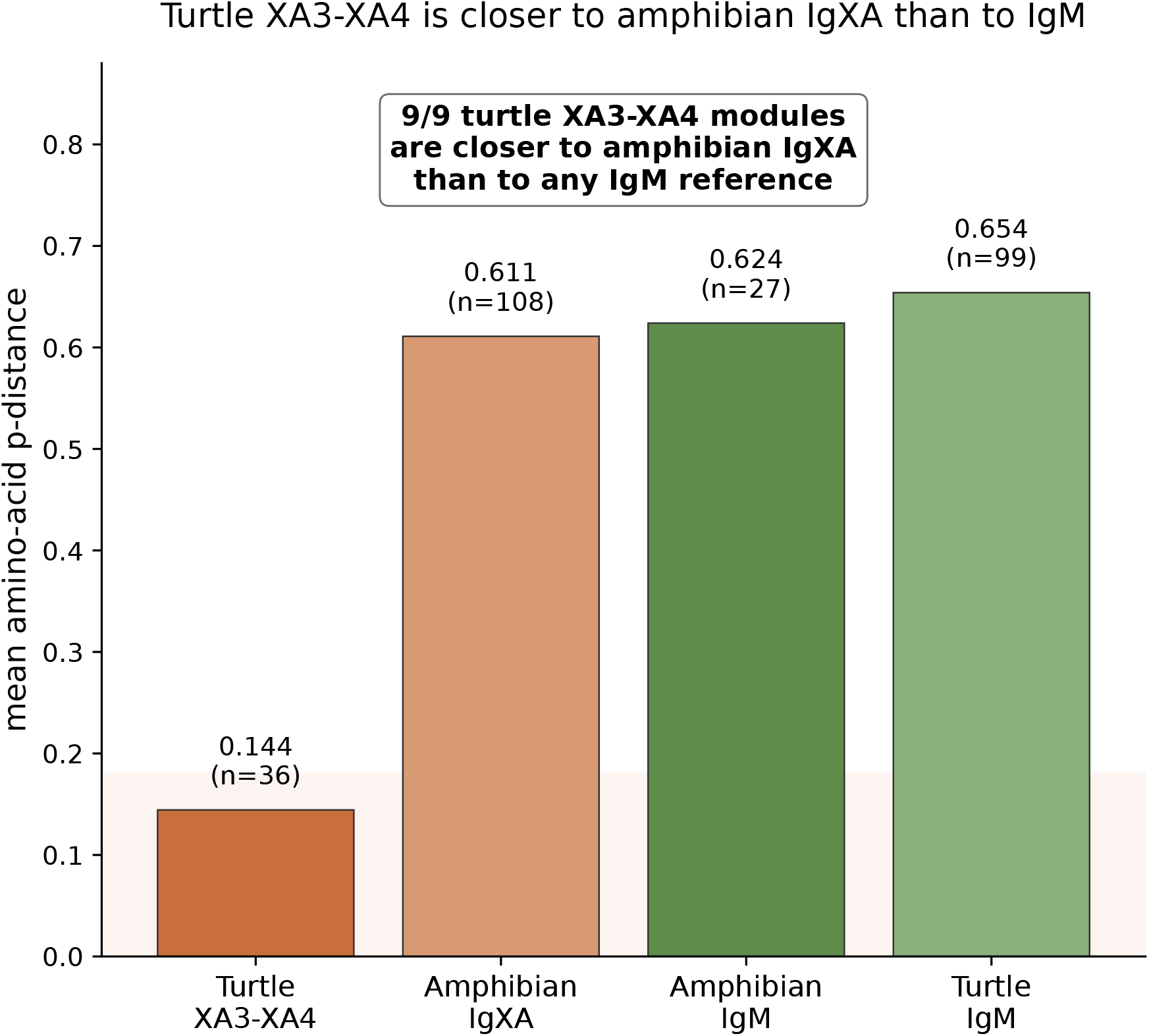
Direct test of turtle XA3–XA4 origin. Paired turtle XA3–XA4 modules were compared with amphibian IgXA3–IgXA4, amphibian IgM3–IgM4, and turtle IgM3–IgM4 references. All turtle XA3–XA4 modules were closer to amphibian IgXA than to any IgM reference, supporting inheritance from the ancestral IgXA module rather than a recent independent IgM duplication in Testudines.

### 3.5 The upstream D scaffold does not behave as a single ancient IgD2 paralogue

At the nucleotide level, the upstream D exons showed patterns inconsistent with a simple model in which the full mixed gene is an intact ancient paralogue of canonical IgD. Mean nucleotide distances between canonical IgD and mixed-locus D exons from the same species were lower than the average mixed–mixed distance for D1, D2, D3, and the D1–D4 concatenation (Fig. 4). For the concatenated D1–D4 scaffold, mean canonical–mixed same-species distance was 0.083, compared with 0.130 for mixed–mixed comparisons and 0.109 for all canonical–mixed comparisons.

**Figure 4:**
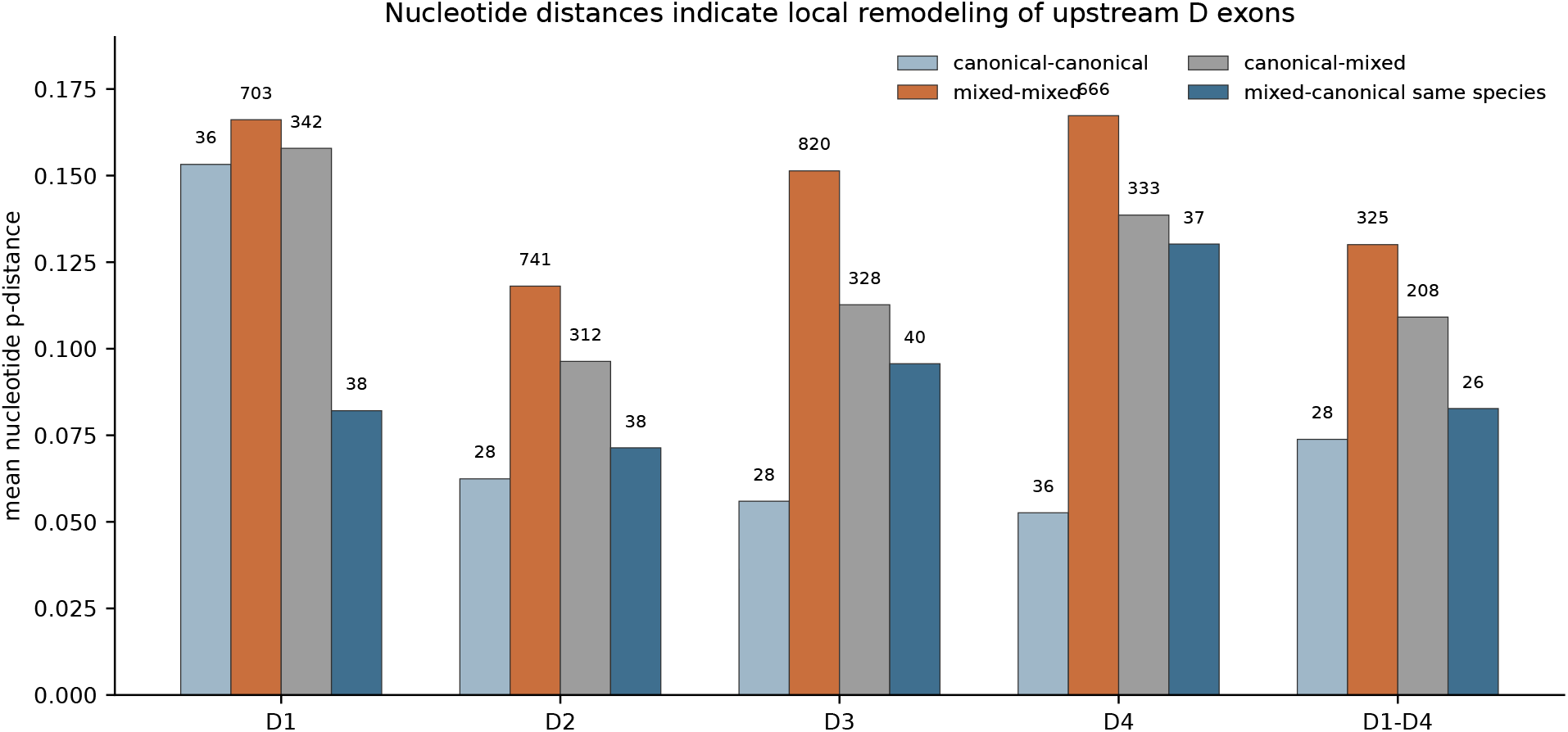
Nucleotide distances for upstream D exons of mixed loci. Mean nucleotide p-distances are shown for canonical IgD comparisons, mixed-locus comparisons, and canonical–mixed comparisons. Same-species canonical–mixed distances are especially low for D1–D3, supporting local remodeling or gene conversion of the upstream D scaffold.

This indicates that the upstream D scaffold frequently retains strong local similarity to the canonical IgD locus of the same species. The effect was strongest for D1–D3. D4 showed a different pattern, with higher canonical–mixed same-species distance (0.130) and fewer local conversion-like cases, suggesting a stronger association with the conserved mixed-locus module.

### 3.6 Conversion-like signal is concentrated in D1–D3

We identified conversion-like cases as mixed-locus D exons for which the nearest canonical IgD exon was closer than any mixed-locus exon from another species. This pattern was observed in 10 D1 cases, 10 D2 cases, 13 D3 cases, but only one D4 case (Supplementary Fig. S3). In the D1–D4 concatenation, six mixed loci showed a local canonical IgD exon closer than any cross-species mixed locus.

Nearest-neighbor counts also supported local dynamics. For individual D1–D3 exons, the nearest neighbor was frequently from the same species. In contrast, D4 more often found its nearest neighbor outside the species, consistent with stronger conservation among mixed loci or reduced local conversion.

The formal tract-permutation test refined this inference (Fig. 5). Among 38 tested D1 mixed exons, 20 had their nearest canonical IgD exon in the same species, three showed significant identical-tract clustering at *q ≤* 0.10, and one met the strict conversion-support criteria. Among 38 tested D2 exons, 14 had a same-species nearest canonical IgD exon, eight showed significant tract clustering, and three met the strict support criteria. In contrast, D3 retained a frequent same-species nearest-canonical signal (18 of 40 tested exons) but no significant clustered tract under this permutation assay. Thus, the upstream D scaffold shows two levels of local IgD affinity: a broad nearest-neighbor signal indicating local derivation or remodeling, and a smaller subset of D1 and D2 exons with long identical nucleotide tracts compatible with recent exon exchange or gene conversion. Most mixed-locus D exons are therefore locally related to canonical IgD but are not near-identical recent replacements.

**Figure 5:**
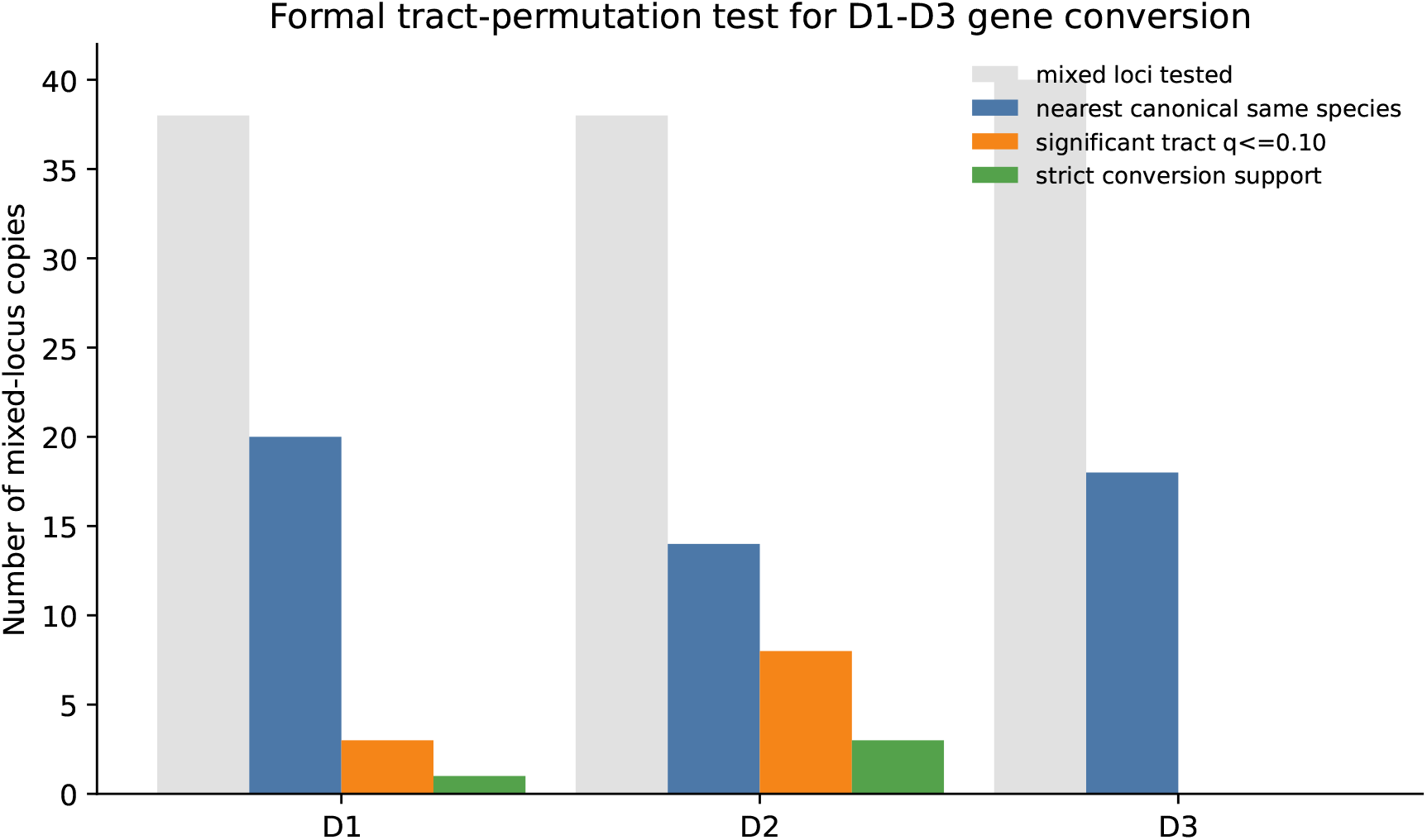
Formal tract-permutation test for D1–D3 local conversion. Bars summarize, for each upstream D exon class, the number of mixed loci tested, the number whose nearest canonical IgD exon is from the same species, the number with significant identical-tract clustering after permutation testing (*q ≤* 0.10), and the number meeting strict conversion-support criteria. D1 and D2 retain formal tract-level evidence for local exchange, whereas D3 shows nearest-canonical local similarity without significant tract clustering in this assay.

### 3.7 IgY genes adjacent to IgD2/XA show inherited copies and local expansion

The same last/*/CHS annotations revealed extensive IgY repetition in the IgD2/XA genomic neighborhood. We identified 391 Y exon calls, 114 local Y blocks, and 76 complete Y1–Y4 IgY blocks. Of the 76 complete IgY blocks, 68 were genomically proximal to an XA exon within 100 kb and all 68 were on the opposite strand from the nearest XA exon. This supports the working genomic model in which repeated IgY genes are interleaved with, or closely adjacent to, complementary-strand IgD2/XA genes.

This proximity was stronger than expected from random local placement within the same contigs (Fig. 6). The randomization test was restricted to 46 complete IgY blocks with high-confidence XA calls on the same contig. All 46 observed blocks were within 100 kb of an XA exon on the opposite strand. In contrast, randomized IgY positions produced a mean of 37.1 opposite-strand proximal blocks, a median of 37, a 95th percentile of 40, and a 99th percentile of 41. No randomization matched the observed value; with the standard plus-one empirical estimate, *p*=5.0 *×* 10*^−^*^5^. The same observed count also exceeded the null distribution when strand was ignored (null mean 41.1, 99th percentile 44, *p*=1.0 *×* 10*^−^*^4^). Thus, the key signal is not merely that individual distances are short, but that complete IgY blocks are consistently embedded in opposite-strand XA-bearing neighborhoods.

**Figure 6:**
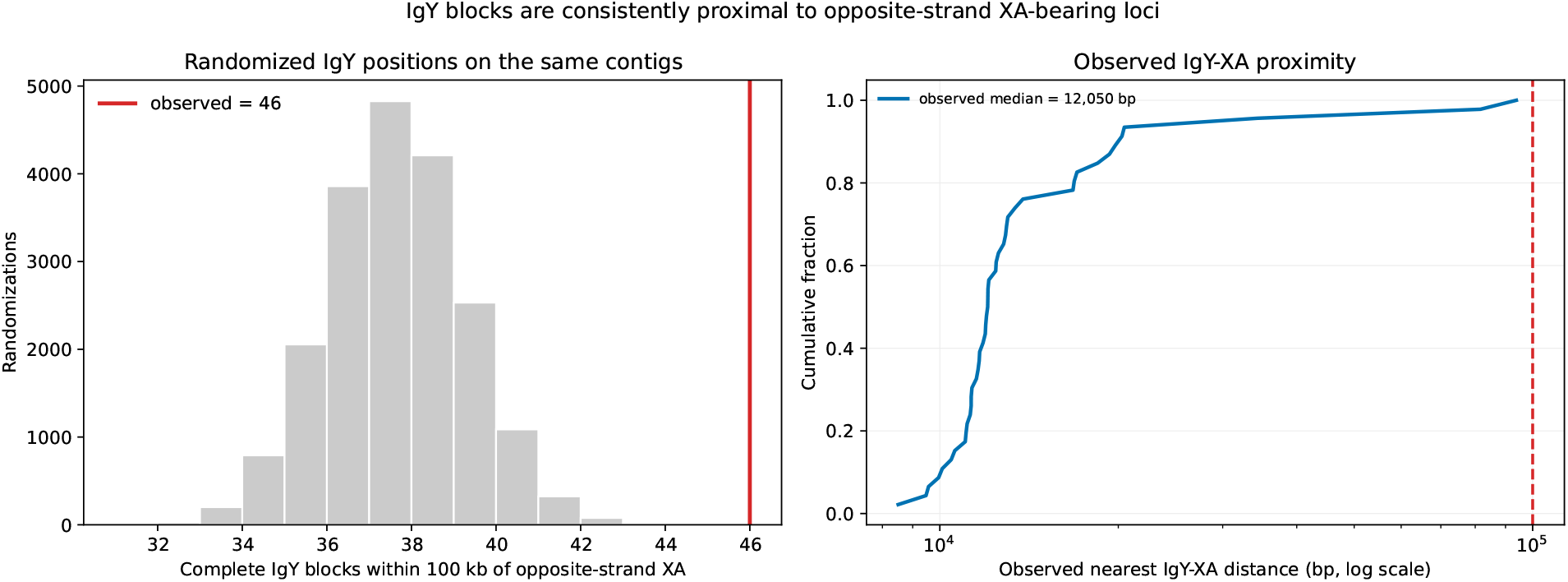
Randomization test of IgY–XA genomic proximity. Complete IgY blocks with high-confidence XA calls on the same contig were compared with randomized IgY positions that preserved contig identity and block length. The primary statistic was the number of complete IgY blocks within 100 kb of an opposite-strand XA exon. The observed value (46 of 46 evaluable blocks) exceeded all 20,000 randomized placements, supporting non-random genomic proximity between repeated IgY blocks and XA-bearing IgD2 loci.

The phylogenetic signal of the complete Y1–Y4 concatenates was not purely species-specific. Same-species pairwise distances were lower on average than different-species distances (mean p-distance 0.171 versus 0.263), and 45 of 76 complete IgY blocks had a nearest neighbor from the same species. However, 31 complete IgY blocks had a nearest neighbor from another species, showing that several IgY copies preserve cross-species orthology or inherited copy structure. This mixed pattern was evident both in nearest-neighbor summaries (Supplementary Fig. S4) and in the complete IgY tree (Supplementary Fig. S5). *Rafetus swinhoei*, *Podocnemis expansa*, *Emydura subglobosa*, and parts of *Mauremys reevesii* formed species-enriched clades with high bootstrap support, consistent with local or tandem duplication. In contrast, other supported clades interleaved copies from different species, including marine turtles and emydid/geoemydid comparisons, supporting inherited IgY copy classes that predate recent species-level duplication. In *Chrysemys picta*, many same-species nearest-neighbor relationships corresponded to the same array represented in two assemblies and should not be counted as independent species-specific expansion without assembly-level curation.

Domain-specific analysis supported a coupled history of the IgY gene (Fig. 7). All six pairs of Y1–Y4 amino-acid distance matrices were positively and significantly correlated (Spearman *ρ* = 0.252–0.730; permutation *p ≤* 0.002). Agreement was strongest for Y1–Y3 (*ρ* = 0.730) and Y3–Y4 (*ρ* = 0.671), whereas Y2–Y4 was the least congruent comparison (*ρ* = 0.252). Depending on the domain pair, 19–34 of 76 genes had exactly the same nearest sequence in both domains, far more structure than expected from unconstrained independent assignments. Thus, IgY domains do not show the pronounced component-level rupture observed between the IgY and D1–D3 trees. The weaker Y2 correlations identify a candidate for nucleotide-level follow-up, but the current amino-acid analysis provides no positive evidence for recurrent IgY domain exchange.

**Figure 7:**
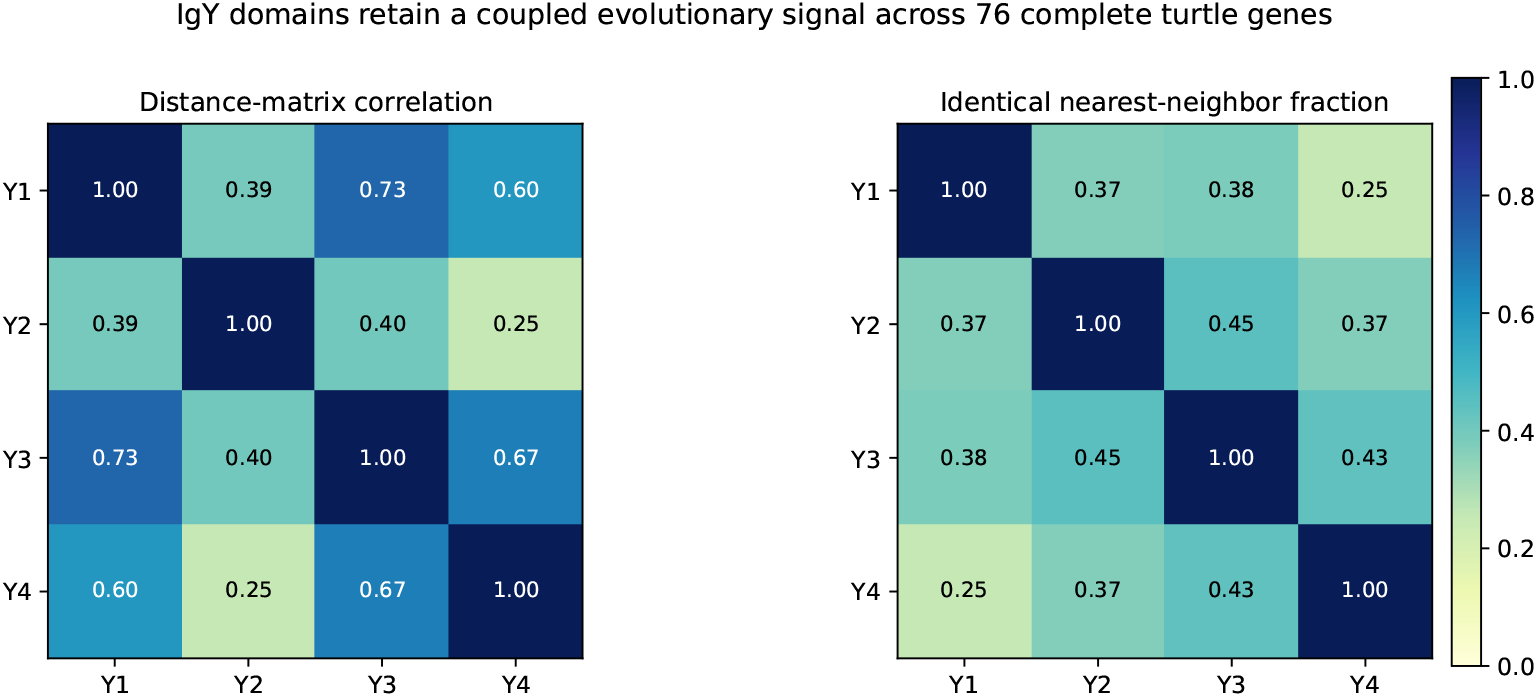
Congruence among the four IgY domains. Left, Spearman correlations between amino-acid p-distance matrices calculated from the same 76 complete IgY genes. All six off-diagonal correlations were significant in 10,000 label permutations. Right, the fraction of genes with an identical nearest sequence in each pair of domain-specific alignments. Y1, Y3, and Y4 retain the strongest coupled signal; Y2 is more weakly resolved but does not show a D-like topological rupture.

Thus, the IgY component records both inherited copy classes and lineage-specific duplication, but it differs from the D scaffold by retaining a more coherent history of the duplicated genomic units. The IgD2/XA locus should therefore be considered within a broader dynamic IgD2–IgY array rather than as an isolated mixed constant-region gene.

### 3.8 Independent marine-turtle arrays replicate paired-module duplication

Reannotation of the *Chelonia mydas* VGP assembly recovered 55 high-confidence IgY exon calls on CM026910.2. These formed 14 local IgY blocks, of which 13 contained complete Y1–Y4 structures. The same interval contained 13 XA-bearing IgD2 blocks, including 13 XA3 and 12 XA4 calls. Conservative one-to-one matching paired all 13 complete IgY blocks with an opposite-strand IgD2/XA block within 100 kb; 12 pairs were within 30 kb and the median interval distance was 5244 bp. Every paired IgY gene was on the reverse strand and every IgD2/XA partner on the forward strand. The array terminates next to a reverse-strand canonical IgD block. Importantly, a thymus transcript assembled from SRR14839285 encoded a VDJ region followed by D1–D4 and XA3–XA4, independently supporting expression of the predicted mixed constant-region architecture.

The independent *Dermochelys coriacea* assembly contained four local IgY blocks on CM019947.2, three of which were complete Y1–Y4 genes. All three complete genes had one-to-one opposite-strand IgD2/XA partners within 100 kb, two within 30 kb, with a median separation of 7445 bp. Self-alignment of both marine-turtle intervals showed repeated off-diagonal matches crossing the array (Fig. 8). The *C. mydas* interval yielded 1752 retained direct and 108 inverted local alignments, whereas the shorter *D. coriacea* interval yielded 140 direct and 31 inverted alignments. These independent genomes therefore replicate physical duplication of larger paired segments rather than independent amplification of IgY alone. Unlike *M. reevesii*, both marine-turtle arrays are predominantly collinear, showing that extensive strand switching is a derived feature of some expanded arrays rather than a requirement for pair duplication.

**Figure 8:**
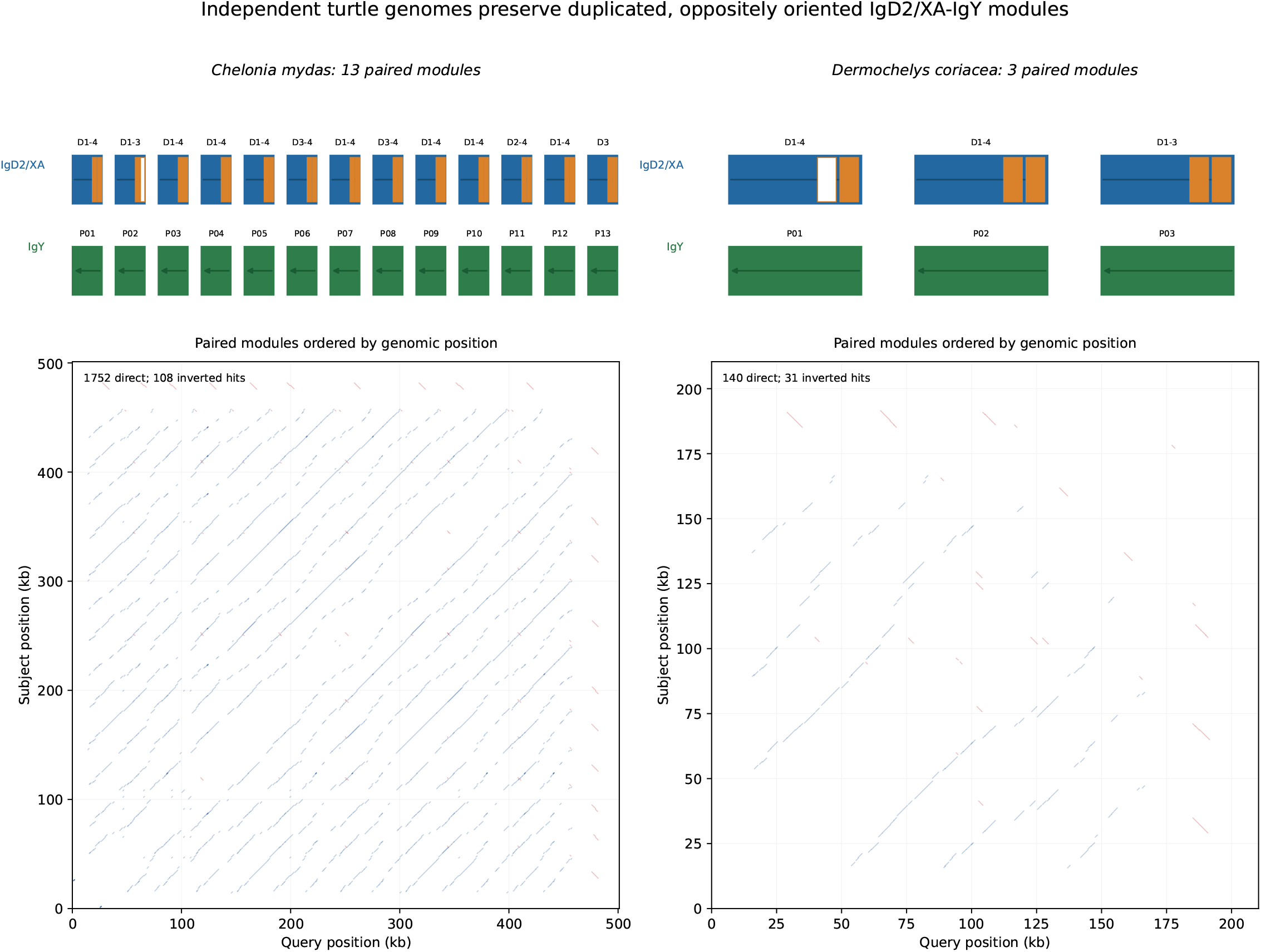
Independent physical validation of paired IgD2/XA–IgY arrays. Uniform-width module strips show complete IgY genes and their one-to-one opposite-strand IgD2/XA partners in *Chelonia mydas* and *Dermochelys coriacea*; white XA boxes mark a missing XA3 or XA4 call. Self-alignments below use physical sequence coordinates. Blue segments are direct and red segments inverted nucleotide matches of at least 700 bp and 70% identity. Repeated off-diagonal diagonals across both loci independently support segmental duplication of paired units.

### 3.9 *Mauremys reevesii* supports duplication of paired IgY–IgD2/XA units

To test whether IgY duplications are accompanied by their IgD2/XA partners, we examined the expanded *Mauremys reevesii* array on contig NC_052635.1 (Fig. 9, Fig. 10; detailed map in Supplementary Fig. S6). Direct curation of the final last/CHS calls recovered additional viable exons that had not been carried into the earlier downstream block table, especially several 4_XA exons. After this curation, 14 opposite-strand IgD2/XA–IgY pairs were retained as analyzable modules. Their physical genomic separation ranged from 4962 to 8352 bp for most pairs, with one more distant putative association at 19,274 bp excluded from the tree analysis because it contained only an isolated IgY exon. The first complete pair, located immediately downstream of the canonical IgD block, contains D1–D4, XA3–XA4, and Y1–Y4 and was labeled Pair01_behind_IgD.

**Figure 9:**
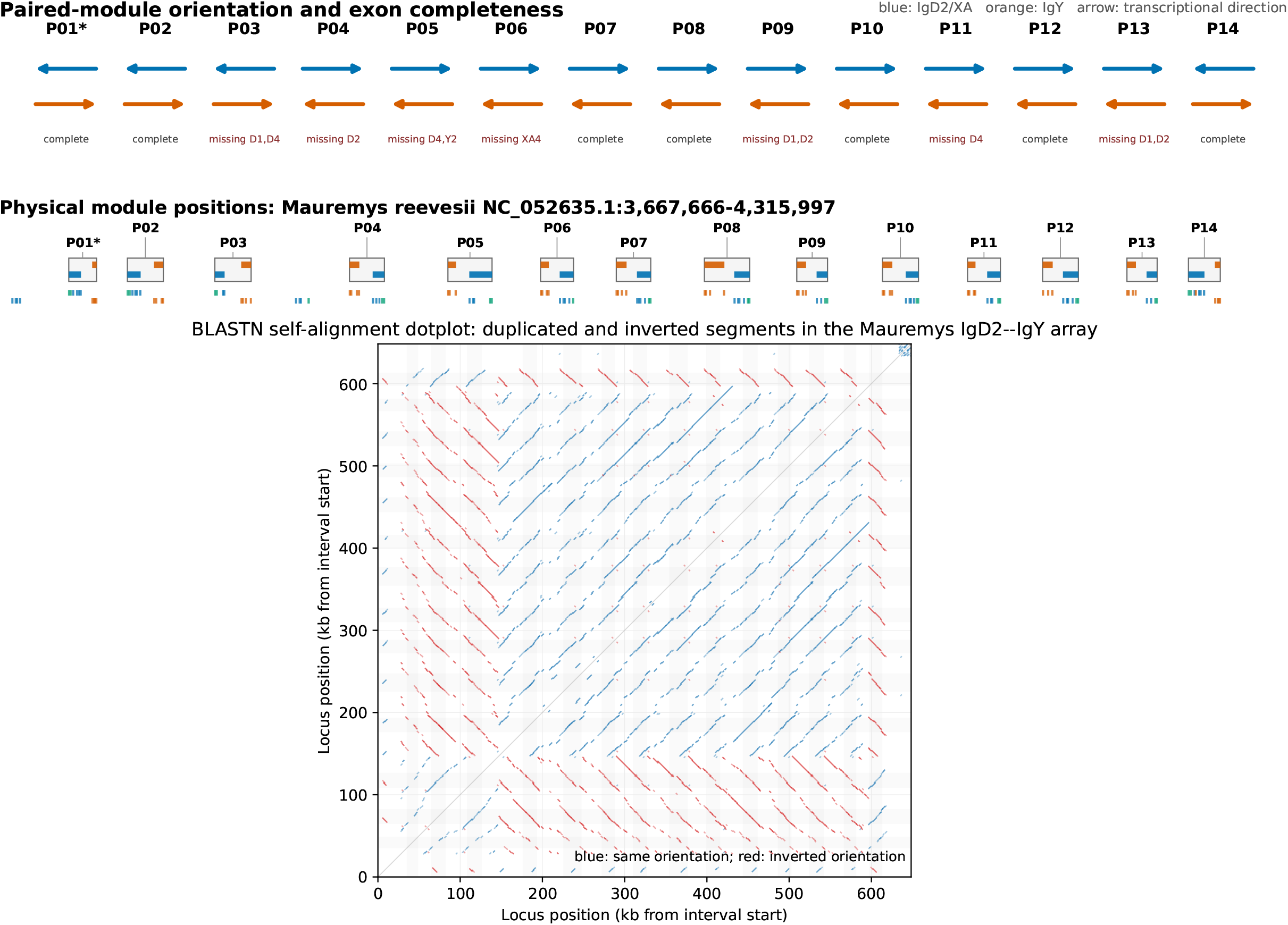
Self-alignment dotplot of the *Mauremys reevesii* IgD2–IgY interval. The upper panel displays P01–P14 at uniform width so that the transcriptional orientation and exon completeness of each IgD2/XA–IgY pair can be read directly; P01 is the first complete pair behind canonical IgD. The middle track restores the physical positions of those modules on NC_052635.1. In the lower self-alignment, blue segments indicate same-orientation alignments and red segments indicate inverted alignments. Repeated off-diagonal signals cross the paired-module positions, supporting segmental duplication followed by inversion or strand switching.

**Figure 10:**
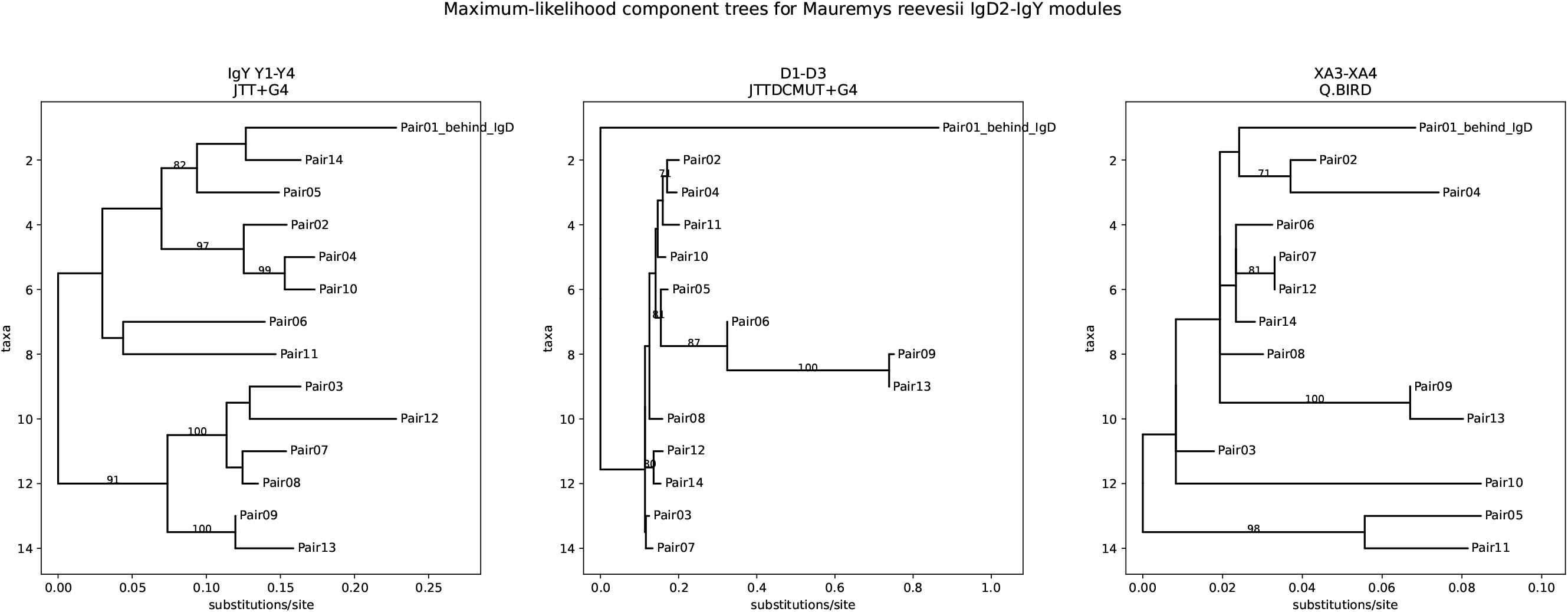
Maximum-likelihood component trees for the *Mauremys reevesii* IgD2–IgY modules. ML trees were reconstructed with IQ-TREE from the same component alignments used for the NJ analyses. Best-fit models selected by BIC are shown above each panel, and ultrafast bootstrap values of at least 70% are labeled. IgY Y1–Y4 retains the clearest repeated-module signal. D1–D3 is discordant with that duplication history, consistent with local D-exon exchange, whereas XA3–XA4 is shorter and partially unresolved.

The architecture is not a simple collinear tandem array. The first three modules have IgD2/XA on the reverse strand and IgY on the forward strand, whereas the central part of the array switches to IgD2/XA on the forward strand and IgY on the reverse strand. The distal region then includes another reverse-strand IgD2/XA paired with forward-strand IgY. This strand alternation indicates that expansion involved inverted segmental duplication or local inversion, rather than straightforward same-orientation tandem duplication. Individual modules also differ in exon content. Several pairs are complete for D1–D4, XA3–XA4, and Y1–Y4, whereas others lack D1/D2, D4, XA4, or one IgY exon. Thus, the duplicated unit is recognizable, but individual copies have undergone secondary exon loss or annotation erosion.

The physical self-alignment supports this interpretation directly. Across the 648,332 bp interval, BLASTN detected 2923 local duplicated segments after filtering, including 1719 same-orientation and 1204 inverted alignments (Fig. 9). The repeated diagonals cross the IgD2/XA–IgY module positions rather than being restricted to isolated exons, indicating segmental duplication of larger local units. Every curated pair overlapped multiple self-alignment hits. The proximal Pair01–Pair03 region and the distal Pair14 region showed especially strong inverted-hit enrichment, whereas many central modules showed more same-orientation hits. This pattern provides physical support for duplication of paired IgD2/XA–IgY units followed by inversion or strand switching.

Validation of the exons added during *Mauremys reevesii* curation separated robust recovered exons from tentative architectural placeholders (Supplementary Table S3). Of 16 rescued or putative exons, 11 had a stop-free ORF in at least one coding phase, 13 had an expected length relative to high-confidence local exons of the same class, and all 16 had nearby splice acceptor and donor motifs within the local boundary window. Eight exons met the strong validation criteria, including rescued D, XA, and Y exons and one putative IgY3 gap candidate. Three rescued canonical IgD D6–D8 exons had stop-free ORFs and nearby splice motifs but no local same-label length reference. Five candidates were retained as questionable because no stop-free ORF was found in the genomic interval, including four rescued exon4_XA calls and one putative IgY3 gap candidate. Therefore, the curated map is supported for the major recovered components, but the questionable calls should be treated as provisional and should not be used as primary phylogenetic evidence.

Module-specific neighbor-joining trees separated the duplication signal from D-exon exchange (Supplementary Fig. S7). The midpoint-rooted full-module tree placed Pair01_behind_IgD as a long basal branch relative to the rest of the array, consistent with the possibility that the first pair behind canonical IgD approximates the proximal ancestral unit. The IgY-only tree retained several high-support relationships consistent with duplication of physical units, including Pair04– Pair10, Pair03–Pair12, and Pair09–Pair13. In contrast, the D1–D3 tree reorganized the array substantially, grouping physically separated pairs such as Pair12–Pair14 and Pair03–Pair04. The XA3–XA4 tree was shallower and less resolved, although some pairs remained strongly grouped, including Pair05–Pair11 and Pair09–Pair13. This discordance supports a mosaic mechanism: IgY often tracks duplication of the physical IgD2–IgY module, D1–D3 can be affected by local exchange or exon replacement, and the short XA3–XA4 component has more limited phylogenetic resolution. Maximum-likelihood analysis supported the same component-level contrast (Fig. 10). The IgY Y1–Y4 alignment was the cleanest ML dataset: ModelFinder selected JTT+G4 [13], no sequence failed the composition test, and the ultrafast bootstrap correlation reached 0.999 [14]. The IgY ML tree recovered strongly supported repeated-module relationships, including Pair04–Pair10 and Pair09–Pair13, and placed Pair01_behind_IgD apart from most internal repeated copies. D1–D3 required the JTTDCMUT+G4 model and showed two sequences with more than 50% gaps, one composition-test failure, and near-zero internal branches; despite final bootstrap convergence, its topology was compressed and discordant with the IgY tree. XA3–XA4 used the Q.BIRD model, included one gap-rich sequence and an identical Pair07–Pair12 pair, and showed several near-zero internal branches. Nevertheless, it retained high-support local groupings such as Pair09–Pair13 and Pair05–Pair11. Thus, the ML trees strengthen the conclusion that IgY carries a clearer physical duplication signal, while D1–D3 and XA3–XA4 are shorter, more erosion-prone components with weaker or discordant phylogenetic resolution.

This arrangement supports duplication of a local inverted IgD2/XA–IgY unit rather than independent amplification of IgY alone. Combined with the nucleotide evidence for local D-exon conversion, the *Mauremys reevesii* array provides a concrete mechanism: IgY is carried by segmental duplication of an IgD2/XA-containing module, while the D exons of IgD2 remain available for exchange or conversion with canonical IgD-derived material.

## 4 Discussion

The turtle IgD2 locus is best explained as a mosaic rather than as a complete ancient IgD paralogue. The XA exons are highly distinct from IgD D exons and form a conserved class across turtles. In the context of the amphibian IgA/X framework, the simplest interpretation is that turtle 3_XA and 4_XA represent the retained XA3–XA4 portion of an older IgXA module. The direct paired-module comparison strengthens this inference: every turtle XA3–XA4 module is closer to amphibian IgXA3–IgXA4 than to IgM references. Therefore, the IgM-like signal in 4_XA should not be read as evidence for an independent turtle IgM duplication. It is better interpreted as the strongest retained IgM-like component of the deeper mosaic history of IgXA; the less decisive XA3 signal is likewise consistent with the intermediate ancestry reconstructed in amphibians [9].

The reconstructed IgM tree provides a useful control for this interpretation. IgM can be reconstructed as a conventional M1–M4 constant-region gene in the same genomes and used as an internal reference for terminal IgM-like signal. Against this background, turtle XA3–XA4 does not behave as a recent turtle IgM duplicate. Instead, the paired XA3–XA4 module is closer to amphibian IgXA than to IgM references, while retaining the expected IgM-related trace inherited through the deeper origin of IgXA itself.

The retention of XA3 and XA4 may also have functional implications. In immunoglobulins with terminal IgM-like or IgA-like constant-region architecture, distal constant domains and associated secretory tailpieces can contribute to polymer formation, J-chain association, and interaction with transport or Fc-like receptor systems. Therefore, the conserved XA3–XA4 terminal module in turtle IgD2 is not only a phylogenetic marker: it may preserve part of an ancestral effector platform. This remains a prediction from exon architecture rather than direct functional evidence, but it provides testable hypotheses for future transcript, secretory-tail, J-chain, multimeric-state, and receptor-binding analyses.

The upstream D exons have a different history. If the mixed gene had evolved as a simple, intact paralogue of the 11-domain IgD, one would expect the upstream D exons of mixed loci to group primarily by mixed-locus orthology across species. Instead, D1–D3 frequently show stronger similarity to canonical IgD exons from the same species. This is a hallmark of local duplication, gene conversion, or recurrent reconstruction from the IgD-like constant-region array. The formal tract-permutation test makes this conclusion more precise: only a minority of D1 and D2 exons behave as recent exchange products with long near-identical nucleotide tracts, whereas most upstream D exons retain a broader, older local relationship to canonical IgD. D3 shows this local IgD affinity but no significant clustered tract in the current assay. D4 is an exception: it exhibits fewer local conversion-like cases and a stronger mixed-locus signal, suggesting that it may have been stabilized earlier with the XA module.

These observations imply that constant-region exon evolution in turtles is modular. The XA3– XA4 module has an ancient origin in the broader IgXA system, whereas the IgD-like scaffold upstream of XA is not a faithful record of that same origin. Instead, it appears to have been repeatedly remodeled in the local heavy-chain locus. This distinction is essential: the presence of XA defines the mixed gene, but the D exons upstream of XA cannot be treated as a single orthologous IgD2 scaffold without accounting for local homogenization.

The IgY repeats extend this modular view beyond IgD2 itself. Most complete IgY blocks are genomically proximal to XA-bearing loci and on the opposite strand, matching the genomic organization observed during annotation. The randomization test shows that this pattern is not explained by random placement of IgY-sized blocks within the annotated contig neighborhoods. Some IgY copies preserve cross-species relationships consistent with inheritance from ancestral turtle arrays, whereas others show strong same-species or same-contig nearest-neighbor signal consistent with recent tandem duplication. The separate Y1–Y4 analyses add an important control: all domain distance matrices are significantly correlated, with the strongest agreement among Y1, Y3, and Y4. Y2 is less congruent, but no IgY domain displays the pronounced phylogenetic rupture observed for the exchange-prone D scaffold. The present evidence therefore supports coupled IgY evolution while leaving isolated or older conversion events formally possible.

The physical result is not restricted to *Mauremys reevesii*. *Chelonia mydas* contains 13 conservatively matched complete pairs, a complete expressed D1–D4/XA3–XA4 thymus transcript, and a dense self-alignment pattern across the array. *Dermochelys coriacea* independently contributes three complete pairs and repeated segmental matches. These marine-turtle arrays preserve a predominantly collinear IgY-reverse/IgD2-forward arrangement, whereas the larger *M. reevesii* array alternates module orientation and varies in exon content. The shared unit is therefore the opposite-strand IgD2/XA–IgY pair; inversion and strand switching are secondary outcomes of array remodeling. Together, the three genomes make independent amplification of IgY alone substantially less plausible and identify duplication of the paired segment as the mechanism that generated local copy number.

These data suggest a mechanistic model in which IgD2 participates in local IgD-derived exchange while carrying neighboring IgY through segmental duplication. In this model, the duplicated unit contains IgY and an oppositely oriented IgD2/XA gene, but the unit can subsequently be inverted, partially eroded, or remodeled. The D exons of IgD2 are not evolutionarily inert: they can be replaced, homogenized, or refreshed by local exchange with canonical IgD or other IgD-like copies. This reconciles the main signals observed here. IgY can preserve the history of duplicated genomic units, D1 and D2 can show a different topology because they continue to participate in D-exon conversion, D3 can retain local IgD similarity after remodeling, and XA3–XA4 can show a shallow or partially discordant signal because it is a short retained module embedded in repeatedly copied local segments.

The physical origin of the array can thus be explained by segmental duplication, inversion, exon loss, and subsequent D-exon remodeling. Its biological significance remains unexplained. In particular, there is no established functional reason why an XA-bearing IgD2 gene should be repeatedly maintained next to an oppositely oriented IgY gene, why both genes should be carried together during expansion, or whether their proximity affects expression, class usage, chromatin organization, or recombination. The persistence of this configuration across divergent turtles makes this unresolved functional association a central biological question raised by the locus.

Several limitations remain. First, most exon calls are computational, although the *C. mydas* thymus transcript now validates expression of one complete mixed architecture; copy-specific long-read transcript evidence is still needed across the arrays. Second, the present analysis focuses on the retained turtle XA3–XA4 module; explicit inclusion of amphibian XA1–XA4 sequences, IgY-derived XA1/XA2 references, and broader reptile outgroups will be needed for a formal phylogeny of the complete IgXA system. Third, the tract-permutation test strengthens the evidence for local exchange in D1 and D2, but nucleotide substitution models and dedicated recombination/conversion methods should still be applied, especially to evaluate the weaker D3 signal. Fourth, the amino-acid domain-congruence test strengthens the inference of coupled IgY evolution but does not prove that exchange never occurs; matched Y1–Y4 nucleotide trees and domain-specific conversion tests are still required, particularly for the less congruent Y2 domain. Fifth, the inferred functional consequences of XA3–XA4 retention require direct validation: secreted isoforms, tailpiece integrity, J-chain association, multimeric state, and receptor binding have not yet been experimentally tested in the turtle IgD2 system. Finally, overlap filtering reduces redundant calls but may hide recent copy-specific variation, and incomplete modules can reflect either biological erosion or assembly/annotation loss.

## 5 Conclusions

Turtle XA-bearing heavy-chain constant-region genes are recurrent IgD2 mixed loci. Turtle 3_XA and 4_XA are not evolutionary IgD D exons; they are best interpreted as retained components of the ancestral IgXA module described from amphibian comparisons, with XA3 retaining an intermediate ancestry and XA4 the clearest IgM-like signal. The terminal module may preserve functional potential for J-chain-associated multimeric forms and receptor-mediated effector interactions. The upstream D scaffold, however, shows local remodeling: a minority of D1 and D2 exons have formal nucleotide tract support for recent exchange with canonical IgD-derived material, most other D exons retain a broader local relationship, D3 lacks significant tract clustering, and D4 is more closely associated with the conserved mixed-locus module. IgY copies preserve both ancestral copy classes and lineage-specific duplications; their four domains retain significantly correlated histories without a D-like rupture. Physical arrays in *Chelonia mydas*, *Dermochelys coriacea*, and *Mauremys reevesii* independently support duplication of opposite-strand IgD2/XA–IgY pairs, followed in some lineages by inversion, exon loss, and local D-exon exchange. Turtle IgD2 should therefore be interpreted as part of an ancestral mosaic IgD2–IgY architecture expanded to generate the present germline array. Although this genomic mechanism is supported, the biological reason for maintaining the unusual association remains unknown.

## Statements and Declarations

### Author contributions

Francisco Gambón-Deza conceived the study, curated the comparative immunoglobulin framework, analyzed the data, prepared the figures, and wrote the manuscript.

### Ethics approval

Not applicable. This study used publicly available genome assemblies and computationally derived annotations and did not involve human participants, clinical samples, live animals, or newly collected biological material.

### Consent to participate

Not applicable.

### Consent to publish

Not applicable.

### Data and code availability

The study used publicly available turtle and amphibian genome assemblies and locally generated constant-region annotations derived from those assemblies. Supplementary Table S1 provides the turtle sampling summary, Supplementary Table S2 identifies every amphibian IgXA3–IgXA4 and IgM3–IgM4 reference, Supplementary Table S4 reports comparative one-to-one IgD2/XA–IgY pairing, and Supplementary Table S5 reports IgY domain congruence. The submission package includes processed exon tables, the reference and analysis FASTA files, alignments, distance summaries, phylogenetic tree files, randomization outputs, figure source files, and analysis scripts. These materials should be deposited in a stable public repository and assigned a persistent identifier before journal publication.

## Acknowledgements

The author thanks colleagues in comparative immunology and immunoglobulin genetics for discussions that helped frame the evolutionary interpretation of reptile heavy-chain loci. The author also thanks the institutions, genome consortia, sequencing centers, database curators, and individual investigators who generated the turtle, amphibian, and reptile genome assemblies and sequence datasets used here and made them publicly available through international repositories, including NCBI/GenBank and related public archives. This study was possible because those primary data were released for community reuse.

