## Supplementary Material for "Turtle IgD2 preserves an ancestral IgXA-derived XA3–XA4 module in a duplicated and locally remodeled IgD2–IgY constant-region array"

Francisco Gambón-Deza

Supplementary Figures S1–S7 and Supplementary Tables S1–S5

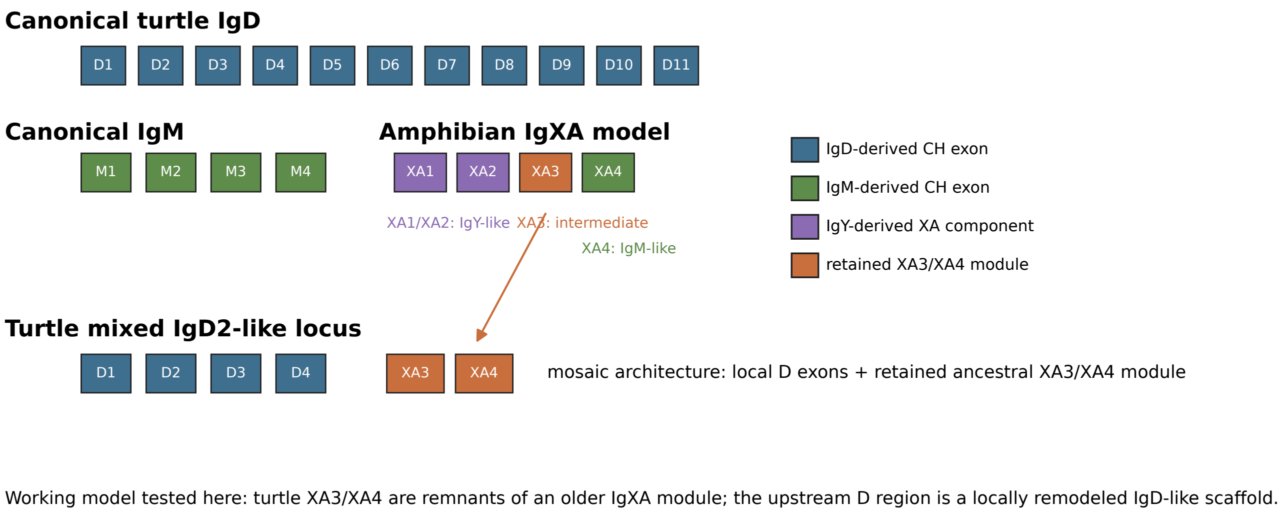


**Supplementary Figure S1.** Working model for the turtle IgD2-like mixed locus. Blue boxes indicate IgD-like D exons and orange boxes indicate the retained XA3–XA4 module.

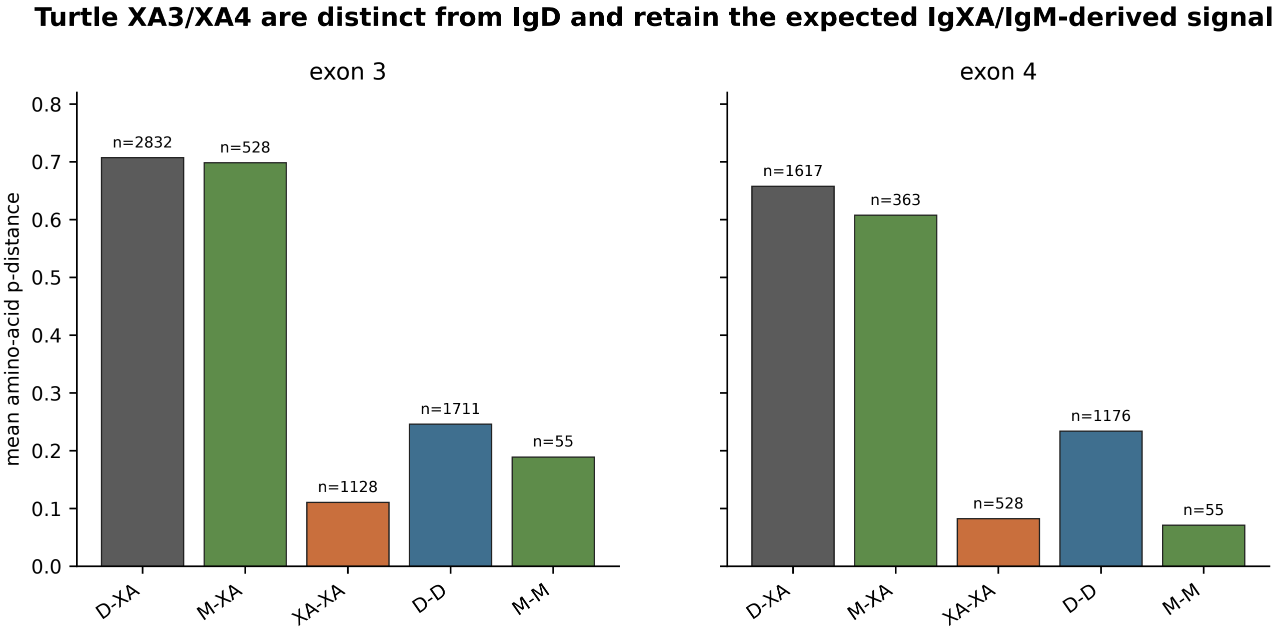


**Supplementary Figure S2.** Amino-acid distances among D, M, and XA exon classes. XA exons form a coherent class distinct from IgD D exons.

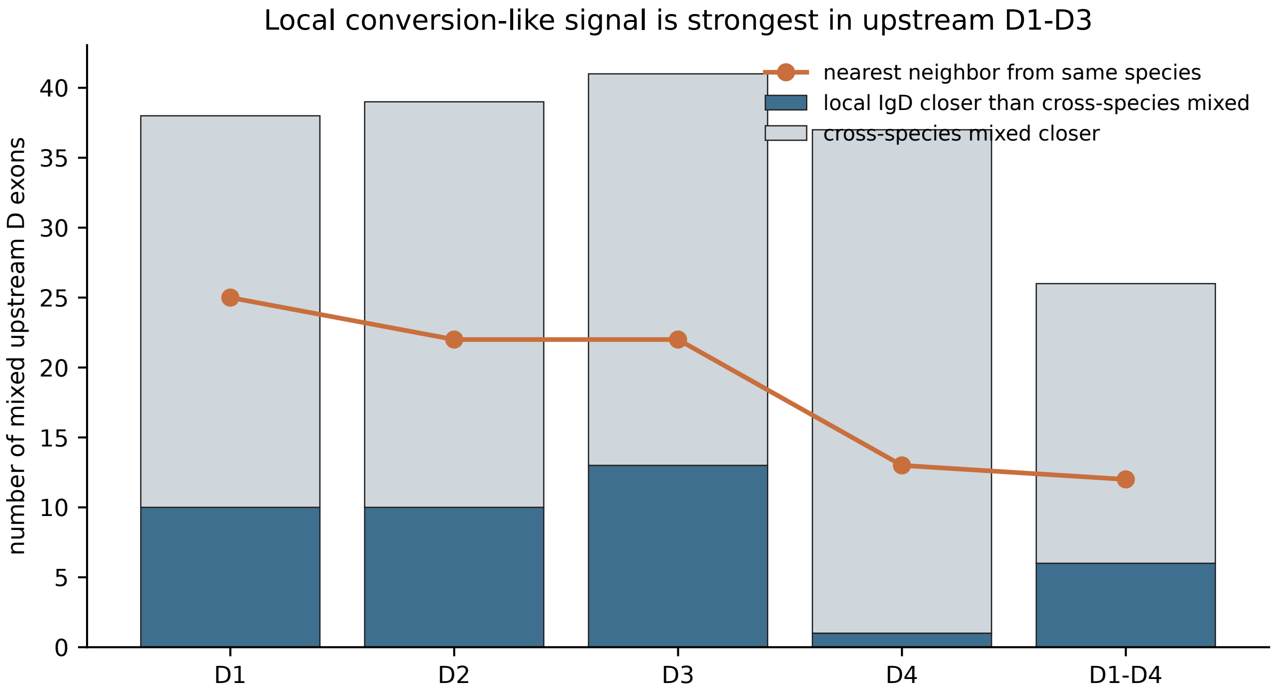


**Supplementary Figure S3.** Local conversion-like signal in upstream D exons of XA-bearing genes.

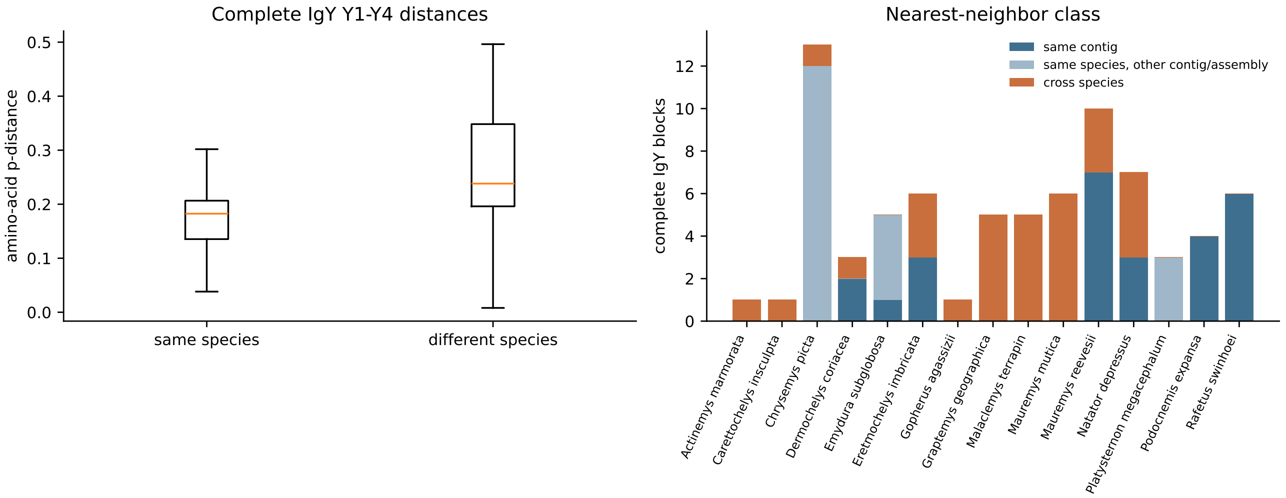


**Supplementary Figure S4.** Phylogenetic and syntenic signal of complete turtle IgY blocks.

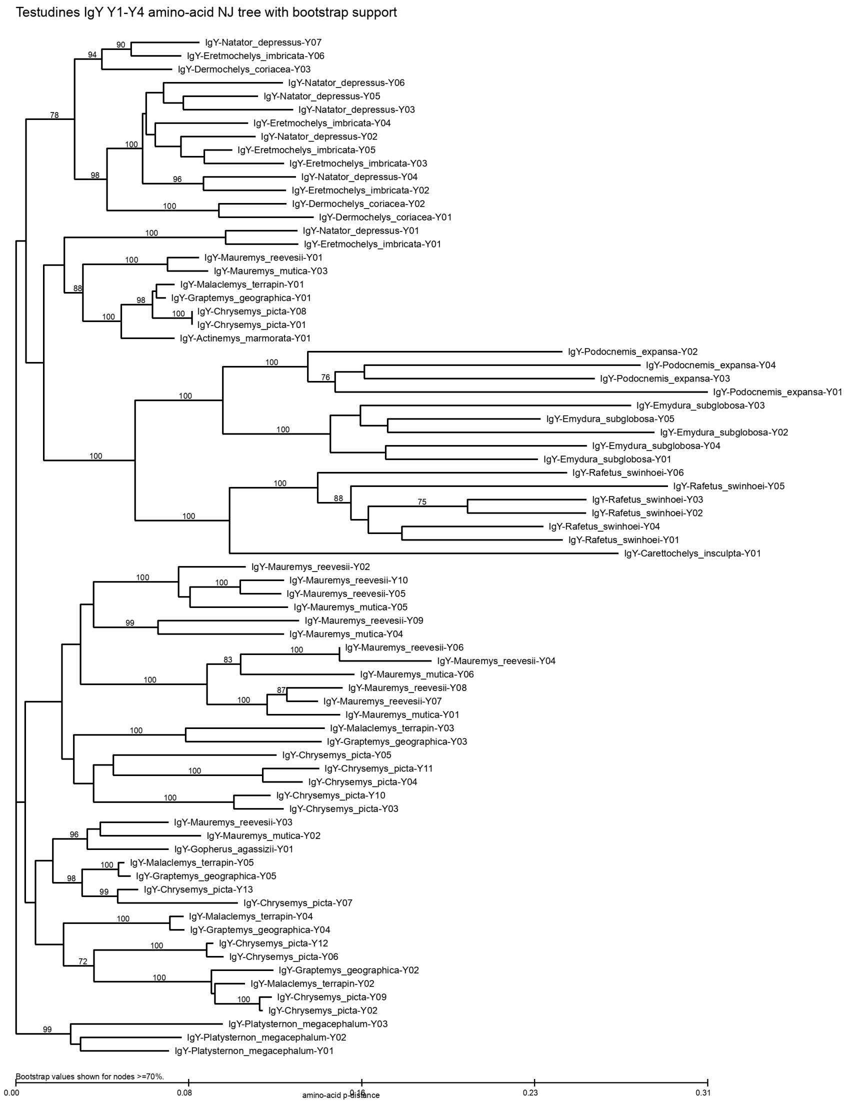


**Supplementary Figure S5.** Neighbor-joining tree of complete turtle IgY Y1–Y4 amino-acid sequences. Node labels report bootstrap support.

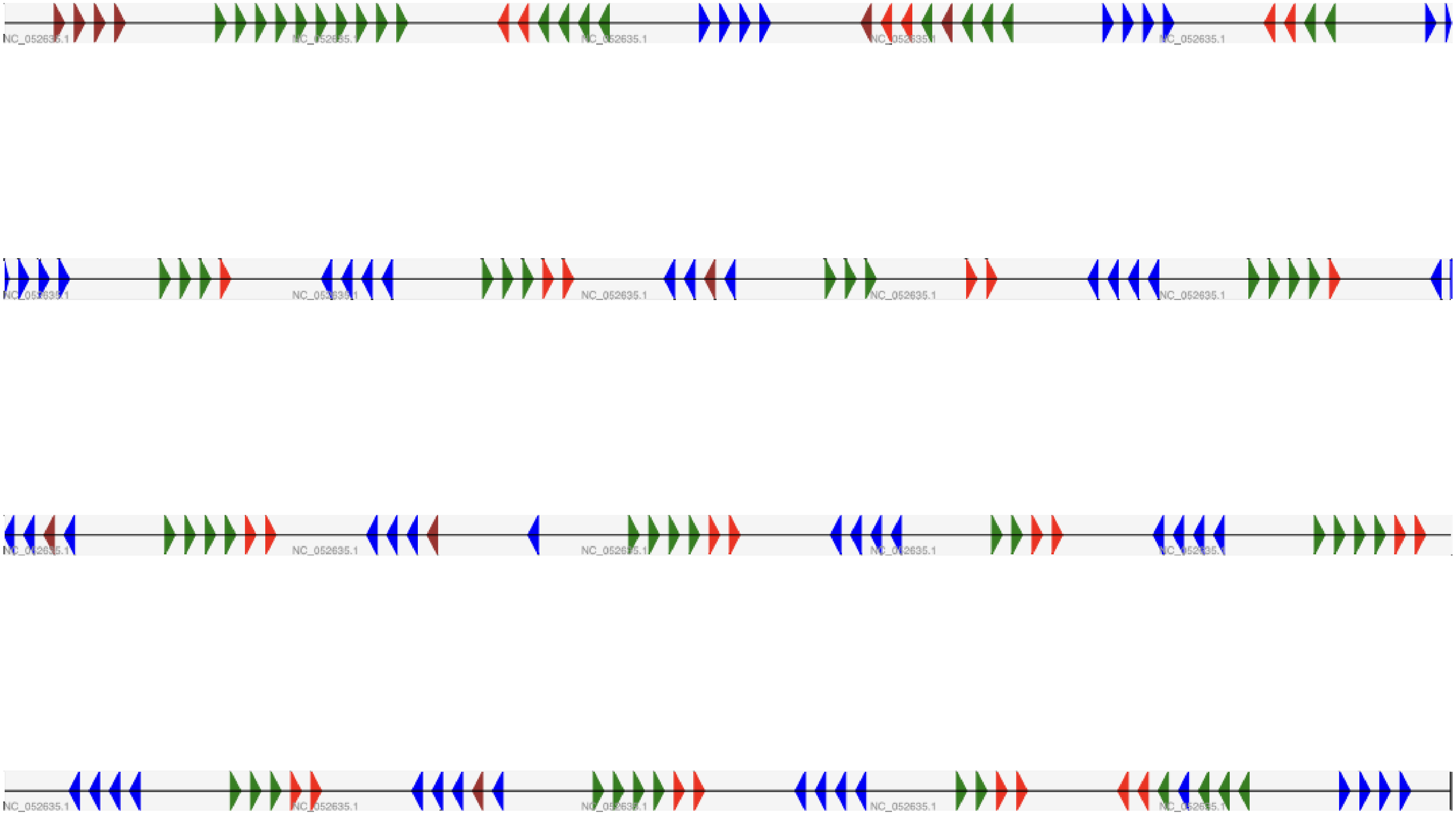


**Supplementary Figure S6.** Curated physical map of the Mauremys reevesii IgD2/XA–IgY array.

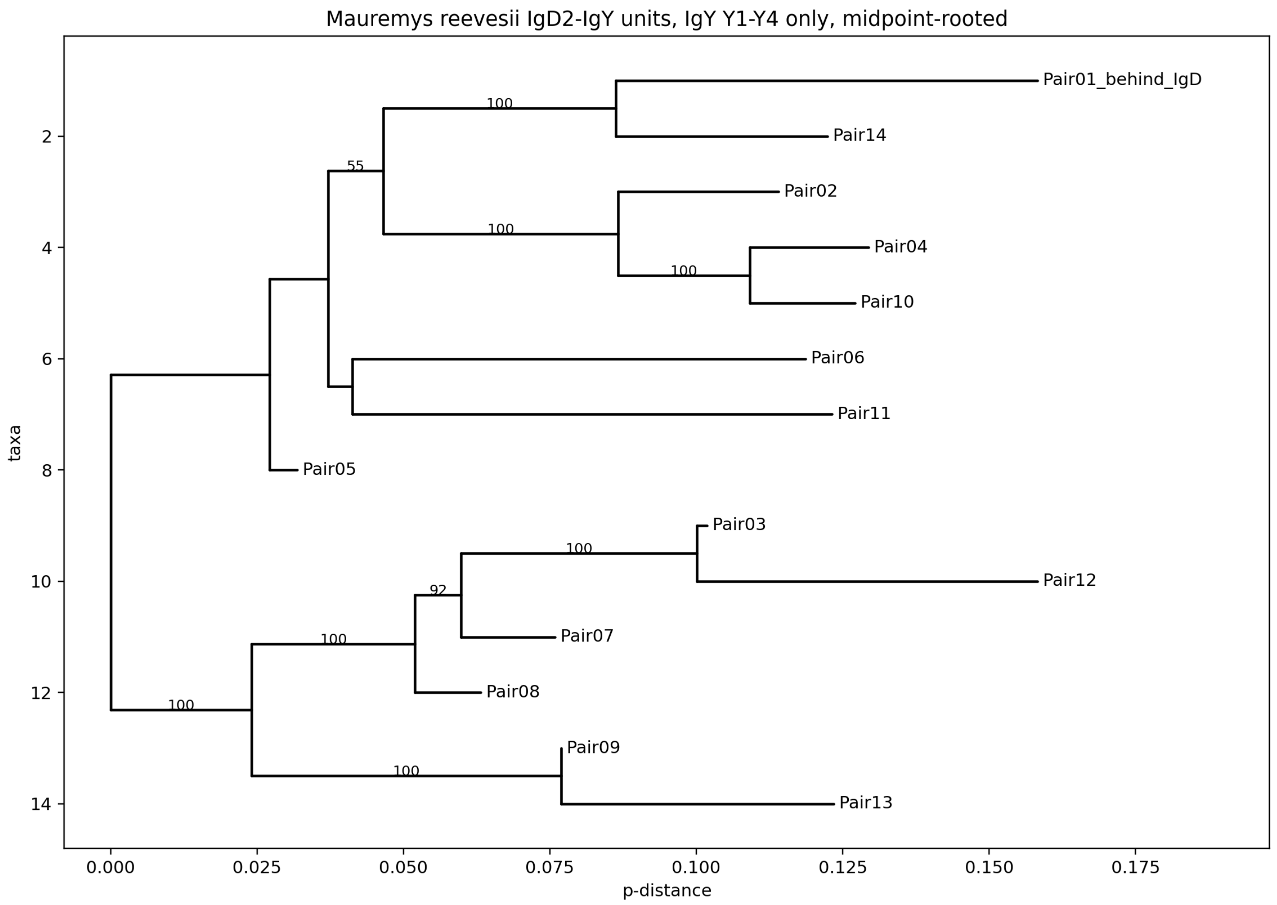


**Supplementary Figure S7A.** Midpoint-rooted neighbor-joining tree of the IgY Y1–Y4 components from paired Mauremys modules.

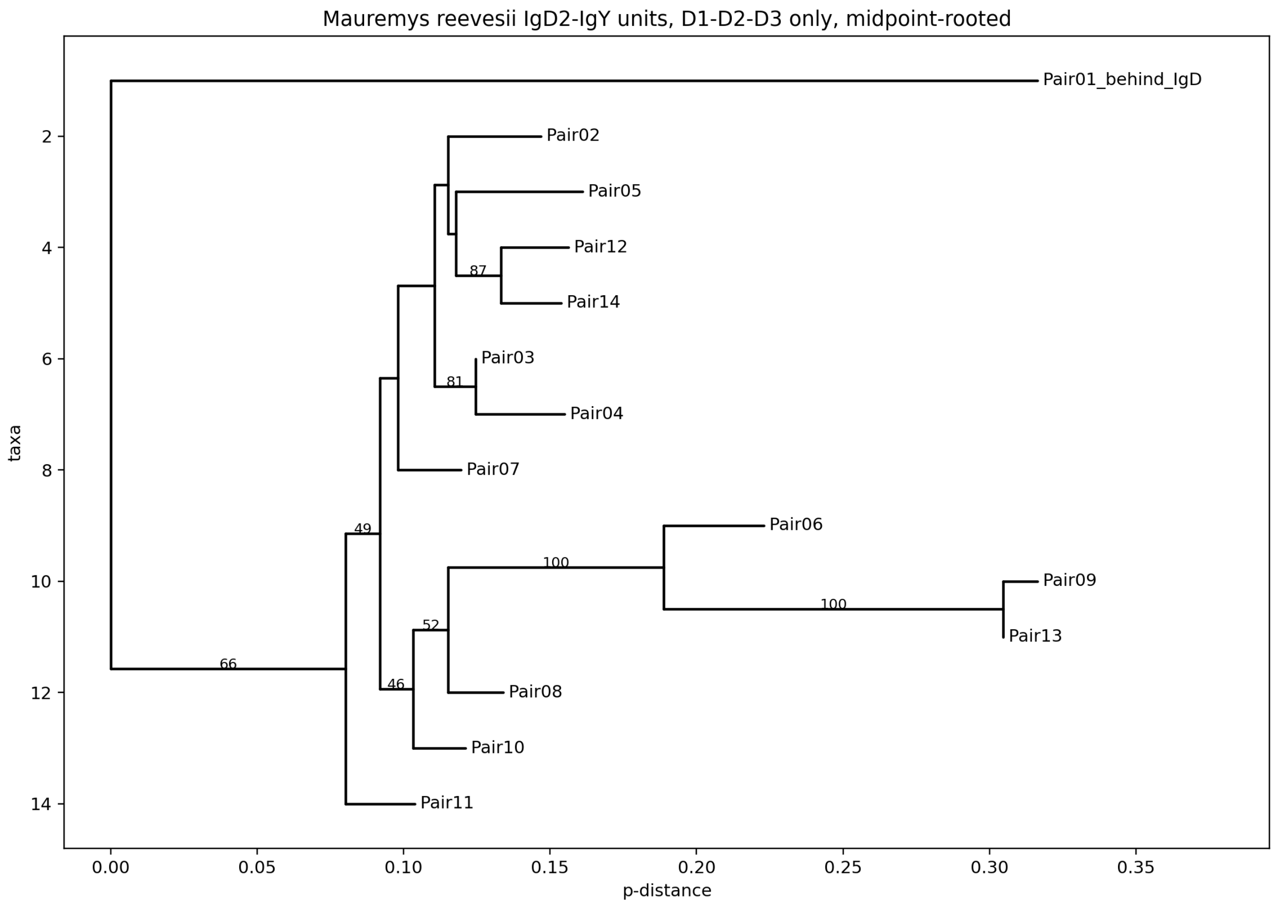


**Supplementary Figure S7B.** Midpoint-rooted neighbor-joining tree of the potentially exchangeable D1–D3 components from paired Mauremys modules.

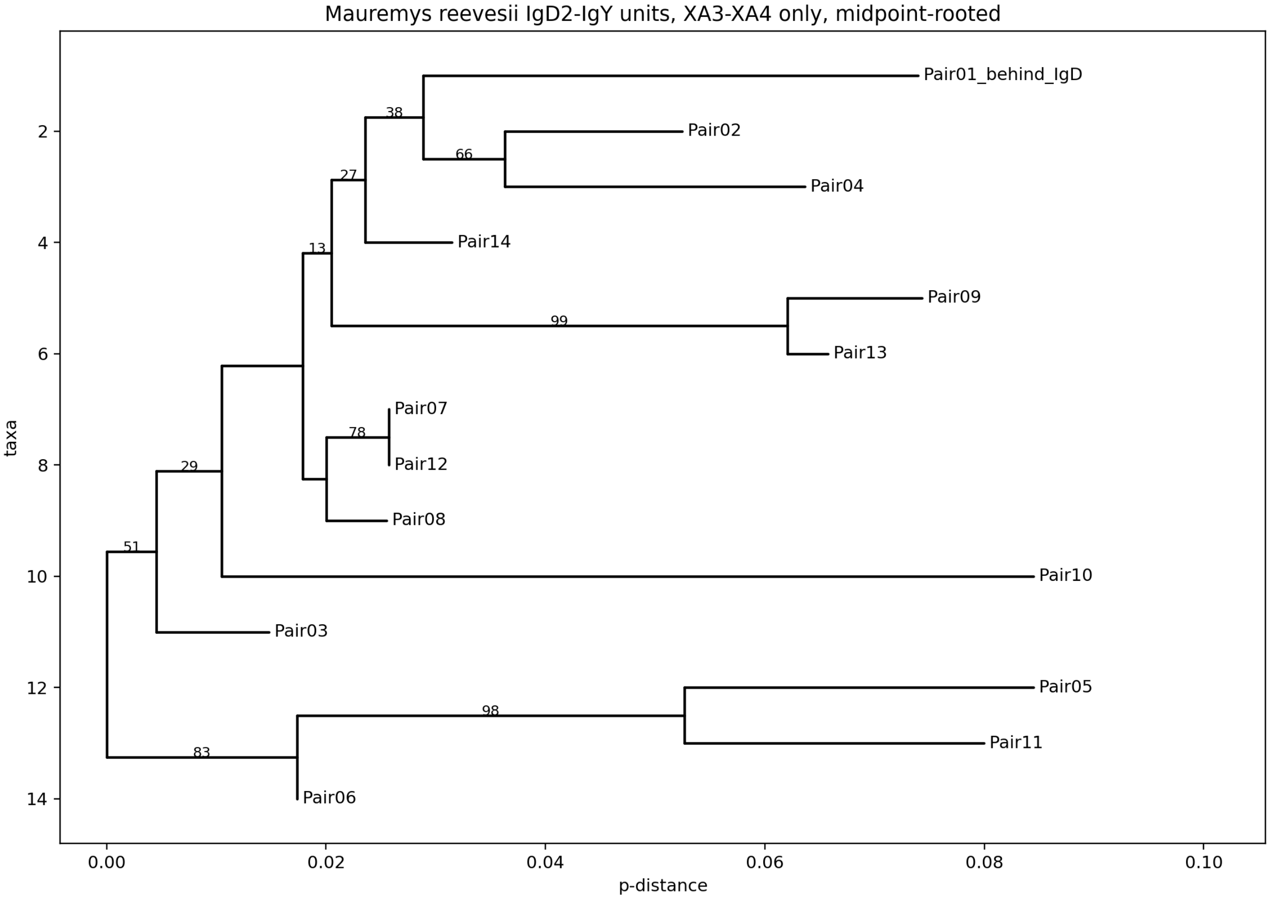


**Supplementary Figure S7C.** Midpoint-rooted neighbor-joining tree of XA3–XA4 components from paired Mauremys modules.

**Supplementary Table S1. Turtle genome sampling and reconstructed constant-region content.**

| Species | Assembly | CHS | IgM contig | Mixed contig | Mixed loci | Complete/near D–XA | Complete IgY |
|---|---|---|---|---|---|---|---|
| Actinemys_marmorata | GCA_009430475.1_Amar_v1_genomic | 47 | ML760385.1 | – | 0 | 0 | 1 |
| Carettochelys_insculpta | GCA_007922185.1_Carettochelys_insculpta-1.0_genomic | 37 | ML683892.1 | – | 0 | 0 | 1 |
| Chelonia_mydas | GCA_015237465.2_rCheMyd1.pri.v2_genomic | 141 | CM026910.2 | CM026910.2 | 13 | 13 | 13 |
| Chrysemys_picta | GCA_037349265.1_rChrPic2_p1.0_genomic | 93 | NW_007281553.1 | JAZIAX010000038.1 | 1 | 1 | 13 |
| Chrysemys_picta | GCF_000241765.3_Chrysemys_picta_bellii-3.0.3_genomic | 82 | NW_007281553.1 | JAZIAX010000038.1 | 1 | 1 | 13 |
| Dermochelys_coriacea | GCA_009764565.4_rDerCor1.pri.v4_genomic | 59 | CM019947.2 | CM019947.2 | 1 | 1 | 3 |
| Emydura_subglobosa | GCA_007922225.1_Emydura_subglobosa-1.0_genomic | 69 | ML680041.1 | ML680041.1 | 1 | 1 | 5 |
| Eretmochelys_imbricata | GCA_030012505.1_ASM3001250v1_genomic | 82 | CM057295.1 | CM057295.1 | 1 | 1 | 6 |
| Gopherus_agassizii | GCA_002896415.1_ASM289641v1_genomic | 39 | PPEB01008883.1 | PPEB01013497.1 | 1 | 1 | 1 |
| Graptemys_geographica | GCA_037349215.1_rGraGeo1_p1.0_genomic | 86 | JBAIFT010000043.1 | JBAIFT010000043.1 | 1 | 1 | 5 |
| Malaclemys_terrapin | GCF_027887155.1_rMalTer1.hap1_genomic | 82 | NC_071516.1 | NC_071516.1 | 1 | 1 | 5 |
| Mauremys_mutica | GCF_020497125.1_ASM2049712v1_genomic | 101 | NC_059084.1 | NC_059084.1 | 1 | 1 | 6 |
| Mauremys_reevesii | GCF_016161935.1_ASM1616193v1_genomic | 165 | NC_052635.1 | NC_052635.1 | 1 | 1 | 10 |
| Natator_depressus | GCA_965152275.1_rNatDep2.hap1_genomic | 93 | OZ223951.1 | – | 0 | 0 | 7 |
| Platysternon_megacephalum | GCA_003942145.1_ASM394214v1_genomic | 65 | QXTE01000558.1 | – | 0 | 0 | 3 |
| Podocnemis_expansa | GCA_007922195.1_Podocnemis_expansa-1.0_genomic | 43 | ML682134.1 | – | 0 | 0 | 4 |
| Rafetus_swinhoei | GCA_019425775.1_ASM1942577v1_genomic | 142 | – | – | 0 | 0 | 6 |
| Trachemys_scripta | GCA_013100865.1_CAS_Tse_1.0_genomic | 23 | CM023067.1 | – | 0 | 0 | 0 |
| Trachemys_scripta | GCF_013100865.1_CAS_Tse_1.0_genomic | 23 | CM023067.1 | – | 0 | 0 | 0 |

CHS, retained high-confidence constant-region exon calls. Blank contig fields indicate that the corresponding representative block was not evaluable under the stated filters.

**Supplementary Table S2A. Amphibian IgXA and IgM references: identity and provenance.**

| Reference | Class | Species | Accession | Strand | Source DOI |
|---|---|---|---|---|---|
| Amphibian_XA_01 | IgXA3-IgXA4 | Leptobrachium leishanense | CM019063.1 | plus | 10.1007/s00251-026-01404-3 |
| Amphibian_XA_02 | IgXA3-IgXA4 | Lithobates catesbeianus | KV956775.1 | minus | 10.1007/s00251-026-01404-3 |
| Amphibian_XA_03 | IgXA3-IgXA4 | Microcaecilia unicolor | NC_044044.1 | plus | 10.1007/s00251-026-01404-3 |
| Amphibian_XA_04 | IgXA3-IgXA4 | Microcaecilia unicolor | NC_044044.1 | plus | 10.1007/s00251-026-01404-3 |
| Amphibian_XA_05 | IgXA3-IgXA4 | Microcaecilia unicolor | NC_044044.1 | plus | 10.1007/s00251-026-01404-3 |
| Amphibian_XA_06 | IgXA3-IgXA4 | Nanorana parkeri | NW_017306493.1 | plus | 10.1007/s00251-026-01404-3 |
| Amphibian_XA_07 | IgXA3-IgXA4 | Pyxicephalus adspersus | CM016421.1 | minus | 10.1007/s00251-026-01404-3 |
| Amphibian_XA_08 | IgXA3-IgXA4 | Rhinatrema bivittatum | NC_042630.1 | minus | 10.1007/s00251-026-01404-3 |
| Amphibian_XA_09 | IgXA3-IgXA4 | Rhinatrema bivittatum | NC_042630.1 | minus | 10.1007/s00251-026-01404-3 |
| Amphibian_XA_10 | IgXA3-IgXA4 | Rhinatrema bivittatum | NC_042630.1 | minus | 10.1007/s00251-026-01404-3 |
| Amphibian_XA_11 | IgXA3-IgXA4 | Rhinella marina | ONZH01018087.1 | minus | 10.1007/s00251-026-01404-3 |
| Amphibian_XA_12 | IgXA3-IgXA4 | Xenopus tropicalis | NC_030677.2 | minus | 10.1007/s00251-026-01404-3 |
| Amphibian_M_01 | IgM3-IgM4 | Leptobrachium leishanense | CM019063.1 | plus | 10.1007/s00251-026-01404-3 |
| Amphibian_M_02 | IgM3-IgM4 | Microcaecilia unicolor | NC_044044.1 | plus | 10.1007/s00251-026-01404-3 |
| Amphibian_M_03 | IgM3-IgM4 | Spea multiplicata | VKOC01000001.1 | plus | 10.1007/s00251-026-01404-3 |

All sequences derive from the amphibian constant-region reconstruction reported by Gambon-Deza (2026).

**Supplementary Table S2B. Amphibian IgXA and IgM references: exon coordinates and annotation support.**

| Reference | E3 start | E3 end | E3 score | E3 aa | E4 start | E4 end | E4 score | E4 aa |
|---|---|---|---|---|---|---|---|---|
| Amphibian_XA_01 | 203484911 | 203485237 | 0.93 | 107 | 203487529 | 203487908 | 0.905 | 107 |
| Amphibian_XA_02 | 19393 | 19722 | 0.984 | 108 | 15788 | 16153 | 0.912 | 114 |
| Amphibian_XA_03 | 29961132 | 29961455 | 0.994 | 106 | 29965029 | 29965381 | 0.995 | 110 |
| Amphibian_XA_04 | 29995625 | 29995948 | 0.993 | 106 | 29998006 | 29998366 | 0.993 | 112 |
| Amphibian_XA_05 | 30030776 | 30031039 | 0.961 | 86 | 30031832 | 30032186 | 0.996 | 110 |
| Amphibian_XA_06 | 222785 | 223114 | 0.99 | 108 | 227961 | 228342 | 0.993 | 124 |
| Amphibian_XA_07 | 35056009 | 35056338 | 0.992 | 108 | 35055020 | 35055370 | 0.975 | 113 |
| Amphibian_XA_08 | 7632389 | 7632712 | 0.998 | 106 | 7630438 | 7630806 | 0.996 | 113 |
| Amphibian_XA_09 | 7606065 | 7606388 | 0.998 | 106 | 7602186 | 7602545 | 0.994 | 99 |
| Amphibian_XA_10 | 7545580 | 7545906 | 0.998 | 107 | 7544888 | 7545274 | 0.984 | 127 |
| Amphibian_XA_11 | 235497 | 235829 | 0.992 | 109 | 233889 | 234221 | 0.984 | 109 |
| Amphibian_XA_12 | 140578886 | 140579209 | 0.955 | 106 | 140577425 | 140577763 | 0.975 | 111 |
| Amphibian_M_01 | 203134812 | 203135129 | 0.905 | 104 | 203136330 | 203136691 | 0.99 | 119 |
| Amphibian_M_02 | 29855277 | 29855576 | 0.972 | 98 | 29857415 | 29857773 | 0.993 | 118 |
| Amphibian_M_03 | 55535712 | 55536050 | 0.951 | 111 | 55536261 | 55536622 | 0.922 | 119 |

**Supplementary Table S3A. Validation of rescued Mauremys reevesii exons: coding and length criteria.**

| Label | Status | Probability | Start | End | Strand | Length | Length status | mod 3 | Stop-free ORF | Phases |
|---|---|---|---|---|---|---|---|---|---|---|
| exon6_D | rescued_from_downstream_table | 0.995501 | 3680320 | 3680639 | plus | 320 | not_available | 2 | yes | 1 |
| exon7_D | rescued_from_downstream_table | 0.713231 | 3681151 | 3681353 | plus | 203 | not_available | 2 | yes | 1 |
| exon8_D | rescued_from_downstream_table | 0.587270 | 3681997 | 3682235 | plus | 239 | not_available | 2 | yes | 1 |
| exon4_XA | rescued_from_downstream_table | 0.999847 | 3776036 | 3776326 | minus | 291 | expected | 0 | yes | 1 |
| putative_exon3_Y | putative_gap_candidate_orf_no_stop | 0.500000 | 3793625 | 3793932 | plus | 308 | expected | 2 | yes | 1 |
| exon1_D | rescued_from_downstream_table | 0.997044 | 3816646 | 3816929 | plus | 284 | expected | 2 | yes | 0,1 |
| exon3_D | rescued_from_downstream_table | 0.999540 | 3819164 | 3819363 | plus | 200 | expected | 2 | yes | 0,1 |
| exon4_XA | rescued_from_downstream_table | 0.994305 | 3822952 | 3823334 | plus | 383 | expected | 2 | no | none |
| exon4_XA | rescued_from_downstream_table | 0.998972 | 3860873 | 3861189 | plus | 317 | expected | 2 | no | none |
| exon4_XA | rescued_from_downstream_table | 0.989724 | 3914897 | 3915249 | plus | 353 | expected | 2 | no | none |
| exon4_XA | rescued_from_downstream_table | 0.999205 | 4129145 | 4129431 | plus | 287 | expected | 2 | yes | 1,2 |
| exon2_Y | rescued_from_downstream_table | 0.690856 | 4157065 | 4157343 | minus | 279 | expected | 0 | yes | 1 |
| exon1_D | rescued_from_downstream_table | 0.998166 | 4163923 | 4164206 | plus | 284 | expected | 2 | yes | 0,1 |
| exon4_XA | rescued_from_downstream_table | 0.995676 | 4170221 | 4170570 | plus | 350 | expected | 2 | no | none |
| putative_exon3_Y | putative_gap_candidate_orf_no_stop | 0.500000 | 4192986 | 4193354 | minus | 369 | expected | 0 | no | none |
| exon4_Y | rescued_from_downstream_table | 0.890722 | 4268577 | 4268876 | minus | 300 | expected | 0 | yes | 1 |

### Supplementary Table S3B. Validation of rescued Mauremys reevesii exons: splice signals and final call.

| Label | Start | End | Acceptor | AG exact | AG within 12 nt | AG delta | Donor | Donor exact | GT within 12 nt | GT delta | Validation |
|---|---|---|---|---|---|---|---|---|---|---|---|
| exon6_D | 3680320 | 3680639 | GA | no | yes | -1 | GT | yes | yes | 0 | coding_splice_supported_no_length_reference |
| exon7_D | 3681151 | 3681353 | GC | no | yes | -1 | GT | yes | yes | 0 | coding_splice_supported_no_length_reference |
| exon8_D | 3681997 | 3682235 | GG | no | yes | -1 | GT | yes | yes | 0 | coding_splice_supported_no_length_reference |
| exon4_XA | 3776036 | 3776326 | GC | no | yes | -2 | TC | no | yes | 1 | strong |
| putative_exon3_Y | 3793625 | 3793932 | GA | no | yes | -1 | GT | yes | yes | 0 | strong |
| exon1_D | 3816646 | 3816929 | GC | no | yes | -1 | GT | yes | yes | 0 | strong |
| exon3_D | 3819164 | 3819363 | GA | no | yes | -1 | GT | yes | yes | 0 | strong |
| exon4_XA | 3822952 | 3823334 | GT | no | yes | -1 | GT | yes | yes | 0 | weak_or_questionable |
| exon4_XA | 3860873 | 3861189 | GC | no | yes | -1 | GT | yes | yes | 0 | weak_or_questionable |
| exon4_XA | 3914897 | 3915249 | GT | no | yes | -1 | GT | yes | yes | 0 | weak_or_questionable |
| exon4_XA | 4129145 | 4129431 | GT | no | yes | -1 | GT | yes | yes | 0 | strong |
| exon2_Y | 4157065 | 4157343 | CC | no | yes | -2 | TG | no | yes | 1 | strong |
| exon1_D | 4163923 | 4164206 | GC | no | yes | -1 | GT | yes | yes | 0 | strong |
| exon4_XA | 4170221 | 4170570 | GT | no | yes | -1 | GT | yes | yes | 0 | weak_or_questionable |
| putative_exon3_Y | 4192986 | 4193354 | GG | no | yes | -2 | TC | no | yes | 1 | weak_or_questionable |
| exon4_Y | 4268577 | 4268876 | AA | no | yes | -2 | TT | no | yes | 1 | strong |

Coordinates and splice motifs are reported in transcript orientation. Delta values give the nearest canonical motif offset from the annotated boundary.

**Supplementary Table S4. Conservative one-to-one pairing of complete IgY and XA-bearing IgD2 blocks.**

| Species | Contig | Complete IgY | IgD2/XA | Pairs within 100 kb | Pairs within 30 kb | Median distance (bp) | Orientation |
|---|---|---|---|---|---|---|---|
| Actinemys_marmorata | ML760385.1 | 1 | 0 | 0 | 0 | – | not evaluable |
| Carettochelys_insculpta | ML683892.1 | 1 | 0 | 0 | 0 | – | not evaluable |
| Chrysemys_picta | JAZIAX010000038.1 | 7 | 7 | 7 | 7 | 6074 | opposite strands |
| Dermochelys_coriacea | CM019947.2 | 3 | 3 | 3 | 2 | 7445 | opposite strands |
| Emydura_subglobosa | ML680041.1 | 4 | 1 | 1 | 0 | 83999 | opposite strands |
| Eretmochelys_imbricata | CM057295.1 | 6 | 5 | 5 | 5 | 5250 | opposite strands |
| Gopherus_agassizii | PPEB01013273.1 | 1 | 0 | 0 | 0 | – | not evaluable |
| Graptemys_geographica | JBAIFT010000043.1 | 5 | 5 | 5 | 5 | 6527 | opposite strands |
| Malaclemys_terrapin | NC_071516.1 | 5 | 5 | 5 | 5 | 6434 | opposite strands |
| Mauremys_mutica | NC_059084.1 | 6 | 7 | 6 | 5 | 6642 | opposite strands |
| Mauremys_reevesii | NC_052635.1 | 10 | 12 | 10 | 8 | 5616 | opposite strands |
| Natator_depressus | OZ223951.1 | 7 | 0 | 0 | 0 | – | not evaluable |
| Platysternon_megacephalum | QXTE01000508.1 | 1 | 0 | 0 | 0 | – | not evaluable |
| Podocnemis_expansa | ML682134.1 | 4 | 0 | 0 | 0 | – | not evaluable |
| Rafetus_swinhoei | CM033433.1 | 6 | 0 | 0 | 0 | – | not evaluable |
| Chelonia_mydas | CM026910.2 | 13 | 13 | 13 | 12 | 5244 | opposite strands |

Not evaluable indicates that no high-confidence XA-bearing block was recovered on the selected complete-IgY contig under the conservative filters.

**Supplementary Table S5. Congruence among IgY Y1–Y4 amino-acid distance matrices.**

| Domain A | Domain B | Complete IgY | Spearman rho | Permutation p | Identical nearest neighbor | Same-species neighbors in both |
|---|---|---|---|---|---|---|
| exon1_Y | exon2_Y | 76 | 0.3924 | 9.999e-05 | 28 | 58 |
| exon1_Y | exon3_Y | 76 | 0.7304 | 9.999e-05 | 29 | 52 |
| exon1_Y | exon4_Y | 76 | 0.6028 | 9.999e-05 | 19 | 46 |
| exon2_Y | exon3_Y | 76 | 0.4013 | 9.999e-05 | 34 | 53 |
| exon2_Y | exon4_Y | 76 | 0.2518 | 0.0019998 | 28 | 50 |
| exon3_Y | exon4_Y | 76 | 0.6706 | 9.999e-05 | 33 | 54 |

Permutation probabilities were estimated from 10,000 label permutations.
